# Systems-Scale Structural Modeling Reveals the Germline Architecture of Immunodominance

**DOI:** 10.64898/2026.06.02.729678

**Authors:** Ellen L. Shrock, Eric Sun, Amrita Dhindsa, Max H. Witwer, Zhou Yu, Danton Ivanochko, I-Hsiu Lee, Brian Coventry, Maria Borras Gonzalez, Jean-Philippe Julien, Stephen J. Elledge, David Baker

**Author notes:** These authors contributed equally (Ellen L. Shrock, Eric Sun). These authors contributed equally (Amrita Dhindsa, Max H. Witwer, Zhou Yu).

## Abstract

The adaptive immune system generates diverse antibodies to protect against infection, yet responses often focus on a limited number of antigenic sites, a phenomenon called immunodominance. Using the SARS-CoV-2 receptor-binding domain as a model, this study combines large-scale antibody sequencing, deep mutational scanning, and AlphaFold 3 structural modeling to investigate the basis of immunodominant epitope selection. The results show that germline-encoded antibody features are a primary driver of immunodominance. Specifically, 76% of RBD-targeting antibodies display conserved gene segment usage and/or germline-encoded HCDR3 motifs. Structural analyses identified recurrent germline-encoded residues within these regions that interact with immunodominant epitopes, and mutating these residues eliminated binding. Mutations in SARS-CoV-2 variants frequently disrupt these interactions and are associated with immune escape; other mutations enable germline-mediated recognition and generate new immunodominant epitopes. These findings indicate that innate features, rather than diverse somatic mutations, determine binding specificity for the large majority of the antibody response and underlie antibody immunodominance.

## Introduction

The immune system relies on a vast and diverse repertoire of antibodies to recognize and neutralize a wide array of pathogens. This diversity is primarily generated through V(D)J recombination, in which variable (V), diversity (D), and joining (J) gene segments are rearranged to form the variable regions of the antibody chains. Heavy chains are formed through recombination of one V, one D, and one J gene segment, whereas light chains (kappa and lambda) are generated through recombination of one V and one J gene segment. Beyond this combinatorial diversity, additional variability arises at the junctions of these gene segments through the insertion and deletion of nucleotides during recombination. As a result, the complementarity-determining regions 3 (CDR3s) of heavy and light chains, which span these junctions, are the most diverse regions of the antibody. In particular, the heavy chain CDR3 (HCDR3) has long been considered the primary determinant of antigen specificity due to its extensive sequence variability^1^. Together, these mechanisms enable the theoretical generation of more than 10^11^ distinct antibody sequences, allowing the immune system to recognize virtually any pathogen^2,3^.

Despite this immense sequence diversity, antibody responses often focus on a limited number of antigenic sites, or epitopes, in a phenomenon known as immunodominance^4,5^. Immunodominance has important consequences for viral evolution and immune protection. Because individuals often mount similar antibody responses to a given pathogen, mutations in immunodominant epitopes can facilitate immune escape at the population level. This dynamic was strikingly illustrated during the COVID-19 pandemic, in which SARS-CoV-2 variants of concern have continued to emerge and circulate within previously immune populations, in part by acquiring mutations that evade shared immunodominant neutralizing antibody responses^6–8^. Immunodominance also poses challenges for vaccine design, as several vaccines elicit antibodies against immunodominant (but not necessarily protective) epitopes rather than subdominant sites targeted by broadly neutralizing antibodies^9–11^.

Although immunodominance is clinically significant, the mechanisms that drive it and the degree to which it shapes the total antibody response remain incompletely understood. Both epitope intrinsic factors, such as epitope accessibility and abundance, as well as B cell-specific factors, such as the frequency and affinity of B cell precursors in the germinal center, have been shown to influence immunodominance, although the mechanisms governing these B cell-specific factors were not explored^12–14^. Two linear immunodominant epitopes were previously observed to interact with germline-encoded antibody regions^15^; however, conformational epitopes, which represent the majority of epitopes, were not explored in this study. Thus, it remains unclear whether germline-encoded interactions serve as a general mechanism underlying immunodominance, and to what degree immunodominance influences the overall antibody response to an antigen.

To answer these questions, a large dataset of structurally informed antibody-antigen interactions is required. Here, we use the SARS-CoV-2 RBD, one of the most extensively mapped antigens in history, as a tractable model system to investigate the fundamental mechanisms underlying antibody immunodominance. This work is not intended as a study of antibody responses to SARS-CoV-2 *per se*. Rather, the extensive datasets generated for SARS-CoV-2 RBD uniquely enable investigation at a scale that is sufficiently comprehensive and systematic to support broadly generalizable conclusions. By leveraging thousands of antibody sequences representing the full human antibody response to SARS-CoV-2 RBD^16^, high-resolution epitope mapping from deep mutational scanning (DMS)^17–19^, and large-scale in silico structural modeling^20^, we move beyond anecdotal observations to provide systematic evidence that germline-encoded binding is a primary driver of antibody immunodominance and, more generally, of epitope selection for the large majority of the antibody response.

## Results

### Antibodies to immunodominant RBD epitopes exhibit biased V gene segment usage

To investigate the mechanisms underlying antibody recognition of immunodominant epitopes, we leveraged the extensive SARS-CoV-2 Spike antibody datasets generated during the COVID-19 pandemic. Unique-lineage antibody sequences were curated from the Coronavirus Antibody Database (CoV-AbDab)^16^ and grouped by the Spike subdomain they target: the N-terminal domain (NTD) (n = 523), wild-type Wuhan-Hu-1 receptor-binding domain (wild-type RBD) (n = 5061), and S2 domain (n = 258). We first asked whether antibodies targeting each Spike subdomain exhibit biased V gene segment usage, which would indicate a role for germline-encoded recognition in shaping immunodominance. To account for natural biases in gene segment usage in the naïve human antibody repertoire, we compared the relative usage of V gene segments among antigen-specific antibodies with their frequencies in the naïve repertoire (see Methods)^21,22^. We observed that specific heavy and/or light V gene segments were significantly enriched among antibodies to each subdomain, and that these patterns differed markedly for RBD, NTD, and S2, suggesting that particular germline sequences are preferentially suited for recognizing each region of Spike (**Figures 1A, S1A, and Supplementary Data**).

**Figure 1.**
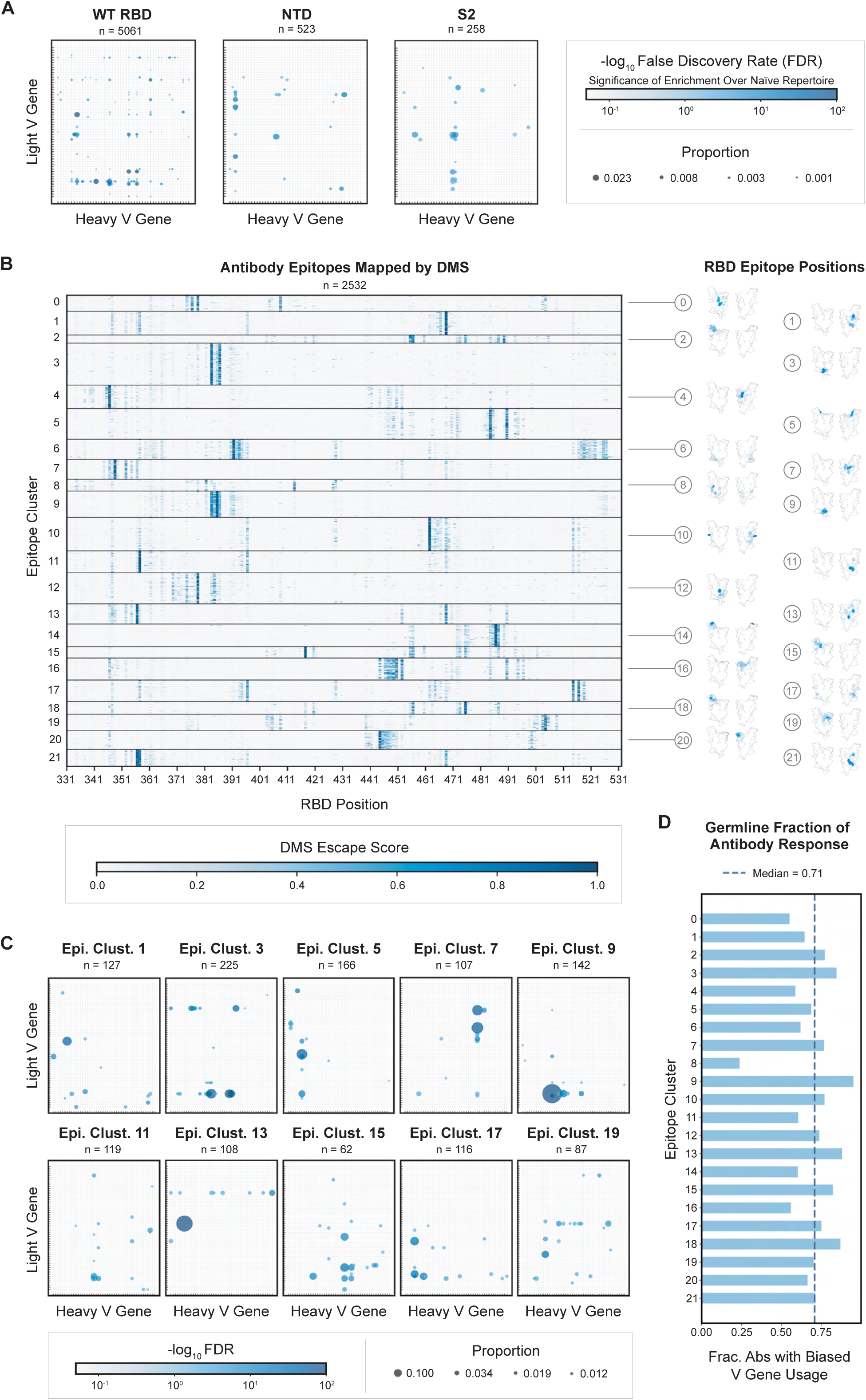
Antibodies to immunodominant RBD epitopes exhibit biased V gene segment usage, suggesting germline-encoded recognition. (**A**) Usage of heavy-light V gene segment pairs among unique-lineage antibodies from the CoV-AbDab targeting the SARS-CoV-2 spike protein RBD (wild-type variant), NTD, and S2 domains, compared to usage in the naïve human antibody repertoire, approximated using 661,780 naïve antibodies from the Observed Antibody Space. Each bubble plot displays the proportion of unique-lineage antibodies that use a specific heavy-light V gene segment pair, represented by the size of the circle. Circle color denotes the statistical significance of enrichment in frequency for that heavy-light V gene segment pair among antibodies to the RBD, NTD, or S2, relative to its frequency in the naïve human antibody repertoire. Enrichment significance was calculated using Fisher’s exact test and corrected for multiple comparisons using the Benjamini-Hochberg False Discovery Rate (FDR) method. Only significantly enriched V gene segment pairs (FDR < 0.05) are shown. (**B**) DMS data for unique-lineage antibodies to the wild-type RBD. Each row corresponds to one unique-lineage monoclonal antibody (in total, data for n = 2,532 antibodies are shown). The values in the heatmap are DMS escape scores, which represent the impact on antibody binding of the mutants at each given position of the RBD (x-axis)^17,18^. Antibodies were clustered by DMS data (see Methods). RBD epitope positions are depicted in blue for each epitope cluster and mapped onto an AF3-predicted wild-type RBD structure, with orientations displaying the inner (left) and outer (right) faces of the RBD. Epitope positions are represented as the proportion of per-residue DMS hotspots relative to the total number of DMS hotspots within an epitope cluster. Colors range from white to marine blue and correspond to a scale of 0 to ≥ 0.25 based on this proportion. Wild-type DMS data were thresholded for escape scores ≥ 0.3 to define DMS-critical residues. (**C**) Bubble plots, as in (A), illustrating heavy-light V gene segment usage for antibodies in each wild-type RBD DMS epitope cluster relative to usage in the naïve human antibody repertoire. (**D**) Proportion of antibodies in each wild-type RBD DMS epitope cluster with significantly enriched heavy V, light V, or paired heavy-light V gene segment usage.

To investigate RBD responses in detail, we analyzed deep mutational scanning (DMS) data for 2,532 unique-lineage antibodies from individuals previously exposed to and/or vaccinated against SARS-CoV-2^17–19^. Each of these antibodies had been assayed individually for binding across a site-saturation mutagenesis library of wild-type RBD, thereby precisely defining the critical epitope residues recognized by the antibody. We clustered the resulting DMS epitope maps into 22 immunodominant epitope clusters (**Figures 1B, S1B, and Supplementary Discussion**), then evaluated V gene segment usage for each cluster. Almost every cluster exhibited striking and distinct biases in V gene segment usage: in some clusters, specific heavy or light V gene segment(s) dominated, while in others, specific pair(s) of heavy and light V gene segments were overrepresented (**Figures 1C, S1C, and S1D**). A median of 71% of antibodies in a given immunodominant epitope cluster exhibited overrepresented V gene segment(s) (**Figure 1D**), suggesting the possibility that most RBD-directed antibodies rely on germline-encoded recognition to engage epitopes.

We investigated whether V gene segment usage was predictive of epitope specificity. Using DMS epitope maps, we performed t-distributed stochastic neighbor embedding (t-SNE) to visualize antibody epitope space in two dimensions. Antibodies containing V gene segments highly enriched among RBD-binding antibodies clustered closely, indicating recognition of similar epitopes (**Figures S2A and S2B**). Consistent with this, averaged DMS epitope maps for antibodies sharing enriched V gene segment pairs revealed strong and well-defined epitope signatures, despite each pairing comprising up to 89 unique-lineage antibodies (**Figure S2C**). Collectively, these findings demonstrate a strong association between germline gene usage and epitope specificity.

### Conserved HCDR3 motifs in antibodies targeting immunodominant RBD epitopes are predominantly germline-encoded

According to the canonical view of antibody recognition, the highly diverse HCDR3 sequences generated by V(D)J recombination are a primary determinant of epitope specificity; therefore, among antibodies to the same immunodominant epitope, we would expect to find conserved HCDR3 motifs. Inspired by work on TCR specificity groups^23,24^, we developed a search algorithm, Sliding Window Investigation of Motifs (SWIM; see Methods), to identify conserved sequence motifs in the HCDR3s of antibodies in each immunodominant epitope cluster (**Figures 2A, 2B, and S3A**). Using this approach, we identified 31 conserved HCDR3 motifs across the dataset (**Figures 2C, 2D, and S3B**). Unexpectedly, the vast majority of these HCDR3 motifs could be traced back to originating from germline-encoded sequences from the D gene segment (79% of HCDR3 motif residues) or J gene segment (3% of HCDR3 motif residues), rather than from non-germline-encoded sequences generated by V(D)J recombination (**Figures 2E and S3B**). Therefore, even when the HCDR3 appears to play a key role in recognizing an immunodominant epitope, it largely does so using germline-encoded sequences. Overall, 76% of the 2,532 antibodies to the wild-type RBD either feature overrepresented V gene segment(s) or a germline-encoded HCDR3 motif, again indicating that the majority of RBD-directed antibodies may engage epitopes through germline-encoded recognition (**Figure 2F and Supplementary Discussion**).

**Figure 2.**
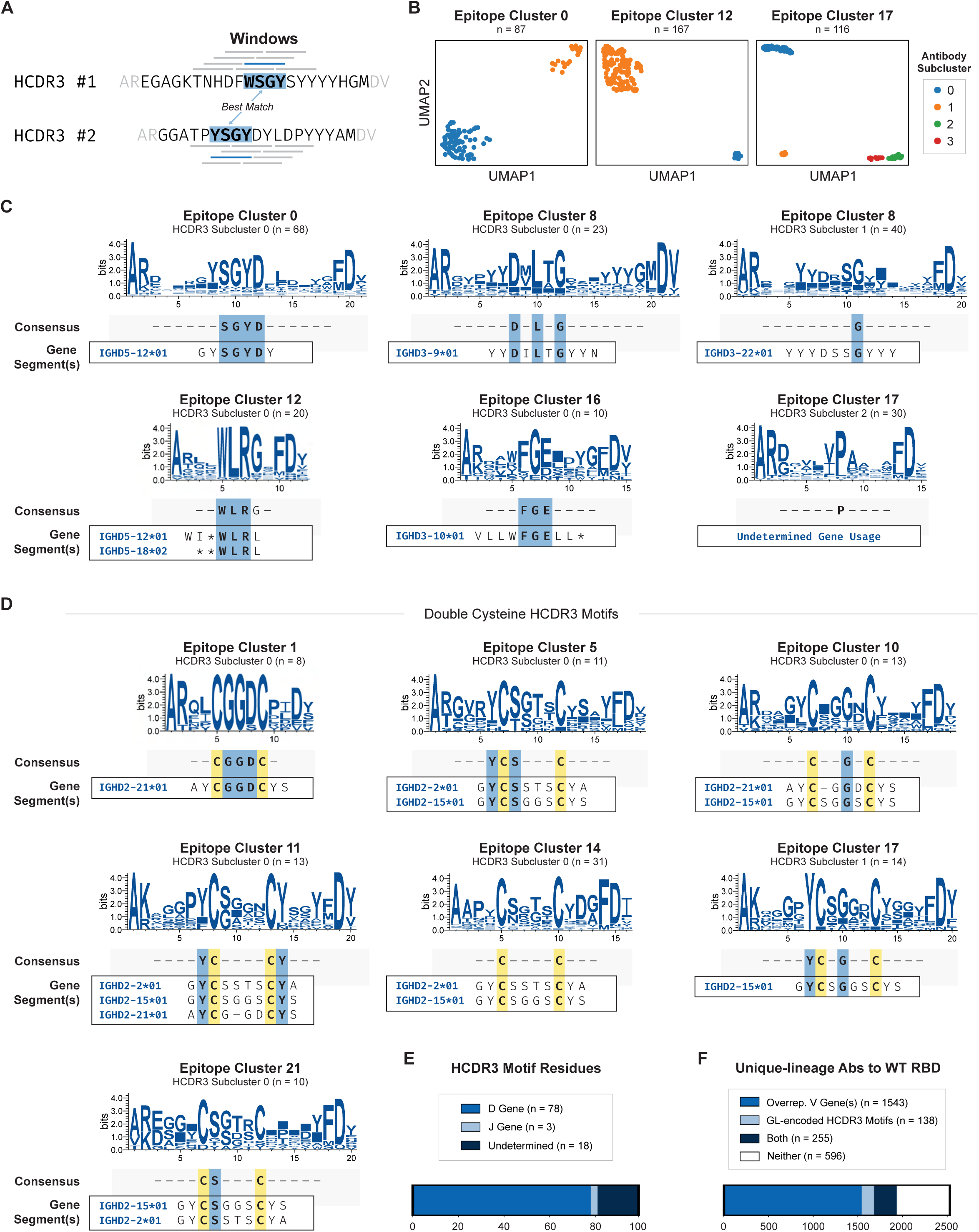
Conserved HCDR3 motifs in antibodies targeting immunodominant RBD epitopes are predominantly germline-encoded. (**A**) The SWIM algorithm samples overlapping windows of four residues from the central region of each HCDR3, determines the similarity between two HCDR3s by measuring the distance between best matching windows, and performs clustering. Distances between amino acids are calculated based on their chemical properties (e.g., hydrophobicity, charge, size). **(B)** UMAP representations illustrate antibody subclusters (in colors) reflecting conserved HCDR3 sequence motifs as identified by SWIM analysis. (**C**) Example HCDR3 subcluster alignments, along with the consensus HCDR3 motif and presumed gene segment(s) from which the motif is derived. HCDR3s in each subcluster were aligned and analyzed using WebLogo^47^. Gaps (solid rectangles) in the logo plots reflect variable HCDR3 lengths. Below each logo plot is the consensus HCDR3 motif, defined as residues with highest (>2) bit value in the third to fourth-to-last position. Below the consensus HCDR3 motif is the likely gene(s) encoding the motif. (**D**) As in (C), but with HCDR3 subclusters that use double cysteine motifs. (**E**) Number of HCDR3 consensus residues that are encoded by a D gene segment, J gene segment, or are undetermined. (**F**) Number of unique-lineage antibodies to the wild-type RBD that employ overrepresented V gene segment(s) (within their corresponding DMS epitope cluster), a germline-encoded HCDR3 motif, both, or neither.

Nine of the HCDR3 motifs, from diverse DMS epitope clusters and collectively found in 5% of the antibodies in the dataset, involved double cysteine residues encoded by IGHD2-2, IGHD2-15, and/or IGHD2-21. AlphaFold 3-prediction of these antibody-antigen complexes (discussed below) revealed that the double cysteines form disulfide bonds, likely stabilizing HCDR3 loop conformation. Thus, these three D gene segments enable the formation of HCDR3 disulfide bonds from their germline-encoded sequences.

### Structural modeling of >1000 antibody-RBD complexes

To directly investigate how antibodies recognize epitopes, we used AlphaFold 3 (AF3) to predict structures for the 2,532 antibodies targeting the wild-type RBD with DMS data and for 1,143 antibodies targeting Omicron BA.5 RBD with DMS data. In addition, we predicted structures for 532 antibodies targeting the N-terminal domain (NTD) and 258 targeting the S2 domain. We modeled each antibody in complex with its respective antigen, and all antibodies represented unique clonal lineages. After filtering for high-confidence predictions (ipTM ≥ 0.8), we obtained structural models for 756 wild-type RBD, 285 BA.5 RBD, 40 NTD, and 5 S2 antibody–antigen complexes (**Figure 3A**).

**Figure 3.**
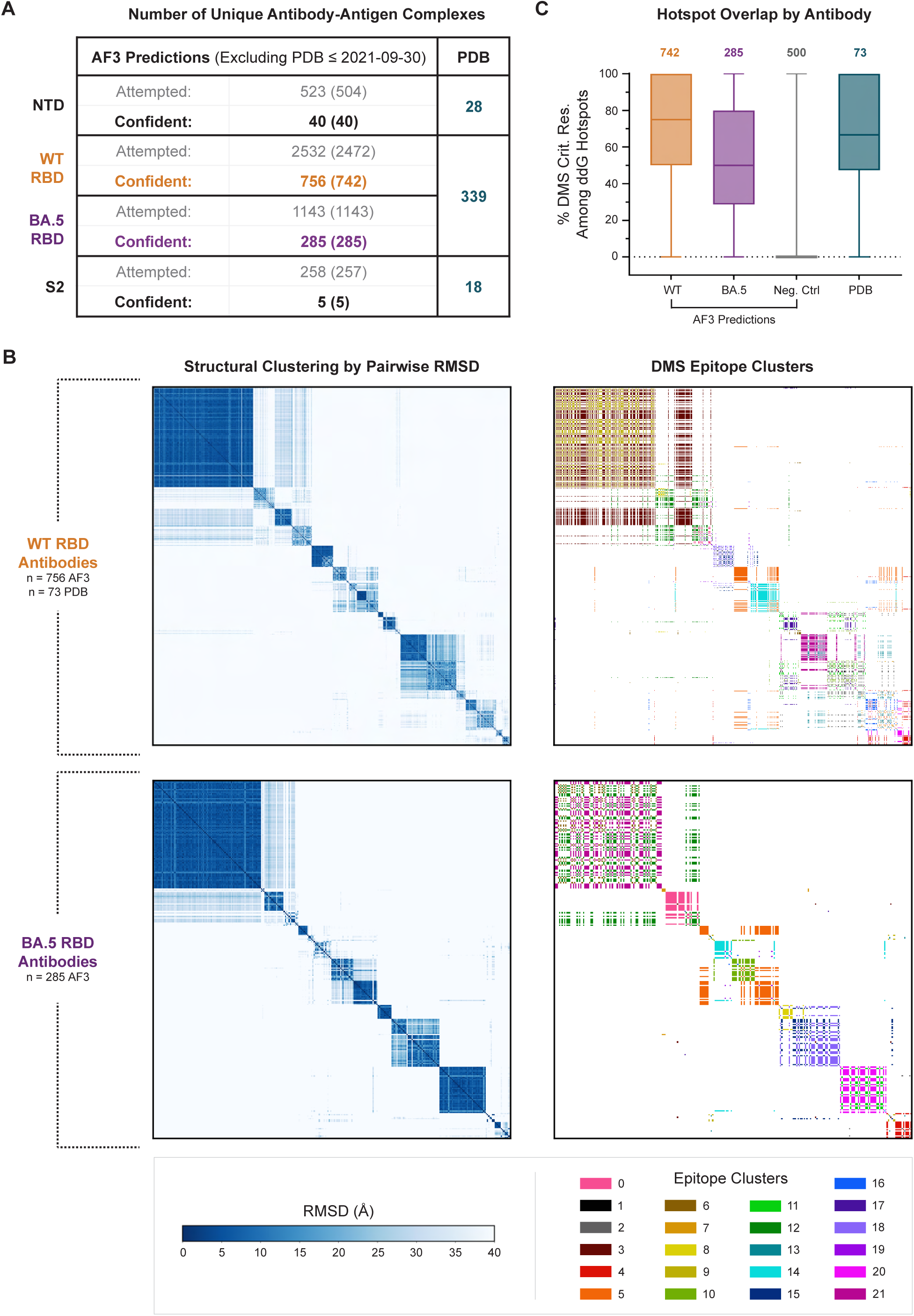
Structural modeling of >1000 antibody-RBD complexes. (**A**) Summary of structural models of antibody-antigen complexes generated using AlphaFold 3 or present in the PDB. The numbers in parentheses in the first column indicate the number of antibodies for which we attempted AF3 predictions excluding the ones that have sequences identical to those of an antibody–antigen complex deposited in the PDB prior to the AF3 training cutoff date of September 30, 2021. (**B**) All-by-all structural clustering of antibody-RBD complexes recapitulates DMS epitope clusters. Left, pairwise RMSDs between corresponding CDR residues in antibody-RBD complexes after aligning on the RBD. When two antibodies share the same DMS epitope cluster, the color is shown. Top row, clustering results for antibody-wild-type RBD structural models with available DMS data. Bottom row, clustering results for antibody-BA.5 RBD structural models with DMS data. (**C**) Agreement between predicted structural models and experimental DMS data. For each predicted or experimentally determined antibody-RBD complex structural model, we plot the percentage of RBD critical residues identified by DMS that were among the residues identified by Rosetta to be interface ddG hotspots.

High-confidence predictions were present for all wild-type and BA.5 RBD DMS epitope clusters, and were more frequent for antibodies binding wild-type (30%) and BA.5 (25%) RBD than for those binding NTD (8%) or S2 (2%), likely reflecting the greater number of experimental antibody-RBD structures in the Protein Data Bank (PDB) used to train AF3 (**Figure 3A**); also, some S2-directed antibodies may target epitopes such as the fusion peptide that are inaccessible in the static S2 structural models predicted by AF3.

To assess the accuracy of the predicted structures, we evaluated their consistency with the experimental DMS data. We first performed all-by-all structural clustering for antibody-wild-type RBD structures and, separately, for antibody-BA.5 RBD structures (see Methods). The resulting structural clusters recapitulated many of the DMS epitope clusters, supporting the reliability of the predicted structural models (**Figures 3B and S4**). We further assessed the reliability of the predicted structures by using Rosetta^25–31^ to identify interface residues predicted to be important for binding, defined as at least 1 kcal/mol change in binding energy (ddG) of the antibody-antigen complex upon in silico mutation ot glycine, hereafter referred to as ddG hotspots. We then calculated, for each antibody-antigen complex, the percentage of DMS-critical epitope residues among the predicted ddG hotspots. As a positive control, in antibody-RBD structures from the PDB with available DMS data (n = 73), a median of 67% of DMS-critical residues overlapped with ddG hotspots. As a negative control, when antibodies were randomly paired to those from a different epitope cluster, we observed a median DMS-ddG overlap of 0%. For AF3-predicted complexes, we observed a median DMS-ddG overlap of 75% for antibodies to wild-type RBD and 50% for antibodies to BA.5 RBD (**Figures 3C and S5**). Altogether, structural clustering and ddG hotspot analyses of the AF3 structures demonstrate strong concordance with the experimental DMS data, supporting their ability to correctly identify key residue-level interactions between antibodies and RBD epitopes.

Of note, not all DMS-critical residues within the RBD necessarily make direct contact with the antibody, as some mutations may disrupt binding indirectly through allosteric effects or by destabilizing the RBD fold. Importantly, the wild-type RBD DMS library was pre-filtered to eliminate mutants that disrupted ACE2 binding, whereas the BA.5 RBD DMS library was not pre-filtered for ACE2 binding and therefore includes mutations that broadly destabilize the RBD. Consistent with this, we observe multiple positions along the RBD that score for nearly all antibodies in the BA.5 RBD DMS dataset (**Figure S1B**), indicative of global structural effects rather than specific epitope contacts. Together, these differences likely account for the reduced DMS-ddG overlap in the BA.5 dataset relative to WT.

### Recurrent germline-encoded interactions are essential for immunodominant epitope recognition

Our observation that gene usage is strongly correlated with epitope specificity suggests that germline-encoded antibody features may be responsible for immunodominant epitope selection. However, an alternative possibility is that these correlations arise indirectly, and that convergent somatic mutations instead determine immunodominant epitope selection. To distinguish between these possibilities, we searched our extensive antibody–RBD structural dataset for structural interactions reused across multiple independent antibodies that bind to the same immunodominant epitope. We reasoned that recurrent structural interactions are more likely to be functional determinants of epitope selection than interactions unique to individual antibodies. If these recurrent interactions are predominantly germline-encoded, this would support a model in which innate features of the antibody repertoire determine which epitopes become immunodominant. Conversely, if they arise primarily from non-germline-encoded residues, this would indicate that convergent somatic mutations play a more dominant role in this regard.

We analyzed the >1,000 predicted structural models of antibody-RBD complexes together with antibody-RBD complexes from the PDB with available DMS data^32^ to identify recurrent structural interactions. We defined a recurrent structural interaction as two or more antibody residues^33^ at Chothia-defined positions that contact an RBD DMS-critical residue or ddG hotspot in at least three unique-lineage antibody-antigen complexes. We grouped these interactions by the V gene segment(s) of the participating heavy and/or light chain(s) and thus identified 1,672 recurrent structural interactions (**Supplementary Data**).

First we investigated whether recurrent structural interactions were predominantly germline-encoded or not. As a reference point, across the structural dataset, antibody residues that contact RBD epitope residues (DMS-critical or ddG hotspots) were evenly split between germline and non-germline origins (47% V gene segment-encoded), and primarily originated from HCDR3 and LCDR3, consistent with the canonical understanding of antibody recognition (**Figures 4A and S6A**). In contrast, antibody residues that were part of highly recurrent interactions (top up-to-five per DMS epitope cluster, collectively observed in over half the dataset) were overwhelmingly (91%) germline-encoded, and primarily originated from HCDR2, LCDR1, and LCDR3 (**Figures 4A and S6A**). Thus, recurrent structural interactions, the putative functional determinants of immunodominant epitope selection, are overwhelmingly germline-encoded.

**Figure 4.**
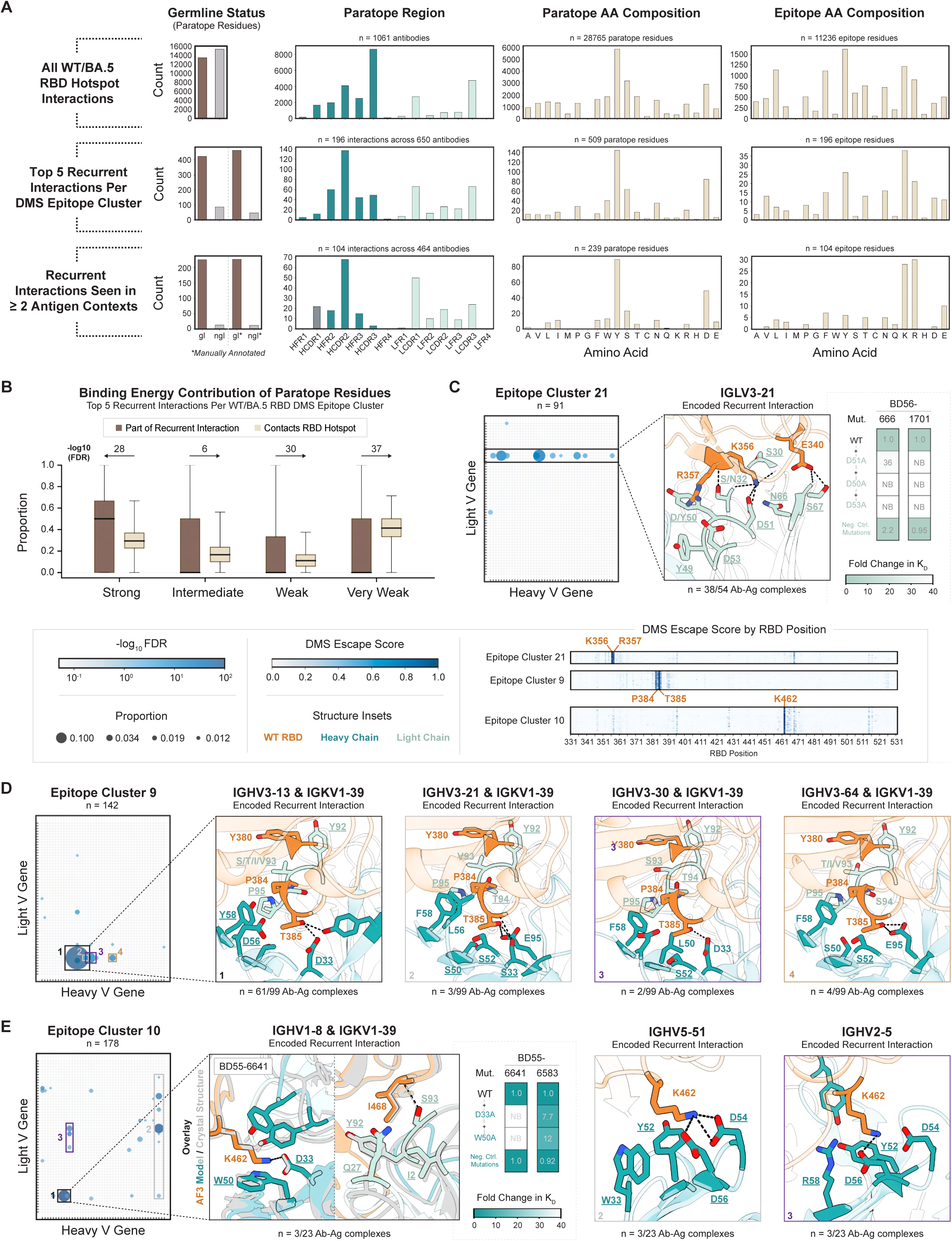
Recurrent antibody-epitope interactions are predominantly germline-encoded. (**A**) Composition of all antibody-RBD structural interactions involving DMS-critical residues or ddG hotspots (top), highly recurrent interactions (top up-to-five per DMS epitope cluster) (middle), and recurrent interactions observed across ≥2 antigen contexts (GRAB motifs) (bottom). The germline status plots indicate whether interacting antibody residues are encoded by V gene segments (gl, brown bar) or not (ngl, grey bar) and, after manual curation, by V, D, or J gene segments (gl*, brown bar) or not (ngl*, grey bar). Antibody residues that are part of highly recurrent interactions are 83% V-encoded and 91% V/D/J-encoded. GRAB motif antibody residues are 95% V/D/J-encoded. Paratope region and amino acid composition plots summarize antibody residues in the interactions indicated by the y-axis; epitope amino acid composition plots summarize the corresponding RBD residues. (**B**) Binding energy contributions of paratope residue subsets: residues involved in highly recurrent interactions (top up-to-five per wild-type and BA.5 RBD DMS epitope cluster; brown) and those contacting RBD DMS-critical or ddG hotspots that are not part of the recurrent interactions (beige). Rosetta-derived per-residue ddG values were classified as strong (ddG < −2 kcal/mol), intermediate (−2 ≤ ddG < −1 kcal/mol), weak (−1 ≤ ddG < −0.5 kcal/mol), or very weak (ddG ≥ −0.5 kcal/mol). For each antibody-RBD complex containing ≥1 highly recurrent interaction (n = 676 structural models representing n = 650 unique antibodies, including both PDB and AF3 structural models) and for each paratope residue subset, the proportion of residues within each ddG category was plotted for both subsets. Boxes indicate the median and interquartile range, while whiskers show the full range. P-values were calculated using the Kruskal-Wallis test followed by bidirectional one-sided Dunn’s post-hoc tests. Z-statistics and directional p-values were computed from overall rank means on the pooled dataset. P-values were adjusted for multiple comparisons using the Benjamini–Hochberg false discovery rate correction applied independently to each tail. The direction of the Dunn’s post-hoc test is indicated by the arrow direction. (**C-E**) Recurrent germline-encoded interactions in DMS epitope clusters 21 (C), 9 (D), and 10 (E). Bubble plots (as in Figure 1C) show heavy-light V gene segment usage of antibodies in the given DMS epitope cluster relative to in the naïve human antibody repertoire. Structure insets depict top recurrent interactions among unique-lineage antibody-antigen complexes from the epitope cluster, with counts annotated. The wild-type RBD is shown in orange; antibody heavy and light chains in teal and pale green, respectively. All antibody side chains that contact the antigen residues are shown as sticks, with dotted lines representing hydrogen bonds. All residues of the identified recurrent interaction are labeled, with germline-encoded residues underlined. DMS data (as in Figure 1B) are shown, depicting the critical residues for each epitope cluster. Fold changes in binding affinity of mutant antibodies relative to the parent antibody, measured by biolayer interferometry, are shown for representative antibodies in epitope clusters 21 (C) and 10 (E) following point mutations of germline-encoded recurrent interactions (see also Figure S6). NB = No detectable binding. In (E), the crystal structure of Fab BD55-6641 bound to wild-type RBD (PDB ID: 11GE) is overlaid with the AF3 model, aligned on the RBD.

Next, we investigated whether antibody residues involved in recurrent interactions are enriched for high binding energy contributions, which would suggest they drive epitope selection. For antibody-RBD complexes exhibiting highly recurrent interactions, we subsetted paratope residues into two categories: those participating in the recurrent interaction(s), and those contacting RBD epitope hotspots (DMS-critical or ddG hotspots) that do not participate in the recurrent interaction(s). Using Rosetta, we calculated per-residue binding energy contributions, classifying them as strong, intermediate, weak, or very weak. We found that residues involved in recurrent interactions were significantly enriched for strong binding energy contributions and correspondingly depleted across all weaker categories relative to other hotspot-contacting residues (**Figure 4B**), suggesting that these recurrent interactions may play a key role in driving epitope selection.

We found that 104 of the 1,672 recurrent interactions appear in the PDB to recognize non-RBD antigens. These motifs, the majority of which are newly reported, are 95% germline-encoded and the residues they target are enriched for lysine and arginine, and to a lesser extent, glutamine. We classify these interactions, which occur within 39 V gene segments, as GRAB motifs because they are observed across multiple antigen contexts, indicating that they are modular structural recognition components within the innate antibody repertoire (**Figure 4A**).

The recurrent structural interactions largely explained the gene segment usage biases and the conserved HCDR3 motifs observed for various DMS epitope clusters (**Figure 1C)**. To illustrate this principle, we highlight examples of DMS epitope clusters where recurrent recognition was mediated predominantly by a single V gene segment (cluster 21), by paired heavy and light V gene segments (cluster 9, cluster 10), by a D gene segment (cluster 0), and by coordinated contributions from V, D, and J gene segments (clusters 14 and 17).

### DMS Epitope Cluster 21: IGLV3-21 encodes a lysine-specific GRAB motif that accommodates an adjacent arginine

For Epitope Cluster 21, IGLV3-21 was strongly enriched (**Figure 4C**). Analysis of the most recurrent structural interactions revealed that in 38 of 54 antibody–RBD structural models for this epitope cluster, a network of germline IGLV3-21-encoded residues (<u>S30</u>, <u>S</u>/N<u>32</u>, <u>Y49</u>, <u>D/Y50</u>, <u>D51</u>, <u>D53</u>, <u>N66</u>, and <u>S67</u>) contact DMS-critical RBD residues K356 and R357, as well E340 (germline-encoded residues underlined; multiple underlined residues at a given position indicate polymorphisms among alleles of the V gene segment; see Supplementary Discussion). The interactions with K356 correspond to a previously described lysine-specific GRAB motif^15^, and IGLV3-21 not only encodes this motif but also accommodates the adjacent R357, providing a structural explanation for its preferential usage (see Supplementary Discussion). Point mutations to IGLV3-21 D50A/D51A consistently abolished antibody binding, whereas control mutations outside of the predicted binding interface had no effect (**Figure 4C, Figure S6B**). These data demonstrate that germline-encoded IGLV3-21 interactions are critical for recognition of this immunodominant epitope.

### DMS Epitope Cluster 9: Coordinate binding by IGHV3-13 and IGKV1-39

Epitope Cluster 9 exhibited strong enrichment for a specific heavy–light V gene segment pair, IGHV3-13 and IGKV1-39, although three additional V gene segments (IGHV3-21, IGHV3-30, and IGHV3-64) were also enriched when paired with IGKV1-39 (**Figure 4D**). Analysis of the most recurrent structural interactions revealed that in 61 of 99 structural models in this epitope cluster, a network of germline-encoded residues from IGHV3-13 (<u>D33</u>, <u>D56</u>, and <u>Y58</u>) and IGKV1-39 (<u>Y92</u>, <u>S</u>/T/I/V<u>93</u>, and <u>P95</u>) bind RBD T385, the most critical residue per DMS data, as well as P384 and Y380. The alternate V gene segments IGHV3-21, IGHV3-30, and IGHV3-64 made similar interactions but used a somatically mutated F58 instead of germline <u>Y58</u>. Together, these data suggest that germline binding by IGHV3-13/IGKV1-39 drives recognition of this immunodominant epitope, with other heavy V gene segments converging via mutation at a key position.

### DMS Epitope Cluster 10: Alternative mechanisms of recognition of K462 by IGHV1-8/IGKV1-39 and IGHV5-51/IGHV2-5

Multiple V genes were enriched in Epitope Cluster 10, including IGHV5-51, IGHV2-5, and the combination of IGHV1-8 and IGKV1-39 (**Figure 4E**). Analysis of the most recurrent structural interactions revealed that in 6 of 23 structural models representing this epitope cluster, IGHV5-51 or IGHV2-5 residues <u>Y52</u>, <u>D54</u>, and <u>D56</u> contact RBD residue K462, the most critical residue per DMS data. In 3 of 23 structural models, IGHV1-8 residues <u>D33</u> and <u>W50</u> engage K462, while IGKV1-39 residues <u>I2</u>, <u>Q27</u>, <u>Y92</u>, and <u>S93</u> contact I468, also a DMS-critical residue for a subset of this cluster.

As there are no PDB structures involving the IGHV1-8/IGKV1-39 recurrent interaction, we obtained a crystal structure of the BD55-6641 Fab in complex with wild-type RBD at 2.76 Å resolution (PDB ID: 11GE), which confirmed the presence of the germline-encoded interactions to K462 and I468. Moreover, point mutant IGHV1-8 D33A ablated or weakened binding to the RBD, whereas control mutations outside of the predicted binding interface had no effect (**Figure 4E, Figure S6C**). We hypothesize that IGHV1-8 is primed to recognize K462 but is insufficient alone; when combined with IGKV1-39 binding to I468, it attains sufficient binding energy to initiate antibody responses. In contrast, the more involved interactions in IGHV5-51 or IGHV2-5 to bind K462 may provide sufficient binding energy to initiate responses alone. Overall, these data provide evidence that the germline-encoded interactions from IGHV5-51/IGHV2-5 or IGHV1-8/IGKV1-39 are important for the selection and recognition of this immunodominant epitope.

### Immunodominance mediated by germline-encoded D segments

For several DMS epitope clusters, the HCDR3 was involved in recurrent structural interactions. However, we observed that the residues involved were germline-encoded by D gene segments, rather than by sequences generated by V(D)J recombination and junctional diversity. Below, we describe three examples.

### DMS Epitope Cluster 0: IGHD5-12 mediates β-strand pairing with the RBD and a GRAB motif-like interaction with K378

Epitope Cluster 0 (**Figure 5A**) was one of a small number of epitope clusters that lacked strong V gene segment biases. Structural modeling revealed a recurrent β-strand pairing between the heavy-chain CDR3 and the RBD, forming an intermolecular β-sheet. Analysis of the most recurrent structural interactions revealed that in 6 of 16 structural models representing this epitope cluster, HCDR3 residues <u>Y100/100A</u>, <u>Y100C/100D</u>, and <u>D100D/100E</u> contact RBD residues K378 and T376, the two most DMS-critical residues for this epitope cluster (numbering differences, e.g. 100/100A, reflect recombination-introduced insertions in some antibodies that change the positions of these residues)^34–36^. SWIM analysis had identified these residues as part of a HCDR3 motif germline-encoded by IGHD5-12 (**Figure 2C**). Point mutations to IGHD5-12 Y100A/D100DA consistently abolished antibody binding, whereas control mutations outside of the predicted binding interface had no effect (**Figure 5A, Figure S6D**). Thus, germline-encoded residues within IGHD5-12 are critical for recognition of this immunodominant epitope. Interestingly, the recurrent interaction targeting K378 resembles several lysine-binding GRAB motifs previously identified in lambda light chains^15^, suggesting a structurally optimized solution for lysine recognition.

**Figure 5.**
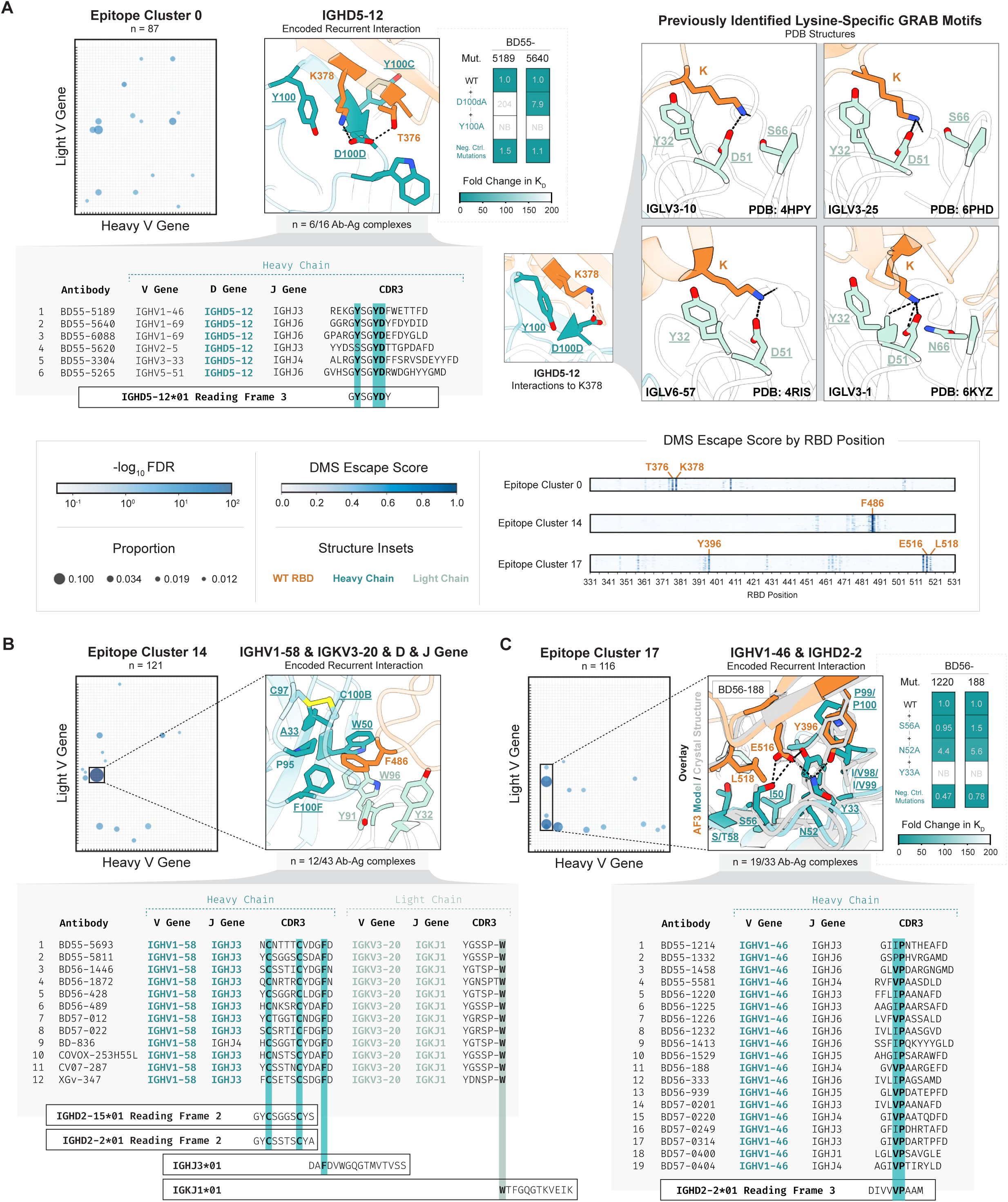
Interactions encoded by germline D and J gene segments, in addition to V gene segments, drive binding to immunodominant SARS-CoV-2 RBD epitopes. (A-C) Recurrent germline-encoded interactions underlying DMS epitope clusters 0 (A), 14 (B), and 17 (C). Bubble plots, as in Figure 1C, illustrate heavy-light V gene segment usage of antibodies in the given DMS epitope cluster, relative to that of the naïve human antibody repertoire. Structure insets, as in Figures 4C-E, depict top recurrent interactions observed among unique-lineage antibody-antigen complexes from the corresponding DMS epitope cluster. DMS data depicting the critical residues for each epitope cluster (defined by escape score) are shown as in Figure 1B. Fold changes in binding affinity of mutant antibodies relative to the parent antibody, measured by biolayer interferometry, are shown for representative antibodies in epitope clusters 0 (A) and 17 (C) following point mutations of their germline-encoded recurrent interactions (see also Figure S6). NB = No detectable binding. In (A), a comparison of the IgHD5-12-encoded interaction to RBD K378 with previously described lysine-specific GRAB motifs found in IgLV3-1, IgLV3-10, IgLV3-25, and IgLV6-57 is shown (right). In (C), the crystal structure of Fab BD56-188 in complex with wild-type RBD (PDB ID: 11GF) is shown as an overlay with the AF3 model of this antibody-RBD complex, as aligned on the RBD. For each epitope cluster, sequence characteristics of antibodies featuring the recurrent interactions in the inset are shown, with conserved features derived from D and J gene segments highlighted and in bold.

### DMS Epitope Cluster 14: IGHV1-58, IGHJ3, IGKV3-20, and IGKJ1 encode an aromatic cage stabilized by a D gene segment-encoded disulfide bond

The heavy–light V gene segment pair IGHV1-58 and IGKV3-20 was strongly enriched in DMS Epitope Cluster 14 (**Figure 5B**)^37–41^. Analysis of the most recurrent structural interactions revealed that in 12 of 43 antibody-RBD complexes, hydrophobic and aromatic residues from heavy (<u>A33</u>, <u>W50</u>, P95, <u>F100F</u>) and light (<u>Y32</u>, <u>Y91</u>, <u>W96</u>) chains form an aromatic cage that encloses the DMS-critical RBD residue F486. With the exception of P95, these residues are germline-encoded by IGHV1-58 and IGKV3-20, as well as IGHJ3 and IGKJ1, and this combination of gene segments is almost exclusively used among antibodies with the recurrent interaction. SWIM analysis identified a conserved HCDR3 motif, CX C, likely encoded by IGHD2-15 and IGHD2-2, that forms a disulfide bond stabilizing HCDR3 conformation and positioning P95 and <u>F100F</u> adjacent to RBD F486. Together, these findings indicate that recognition of the immunodominant epitope in Cluster 14 is mediated by germline-encoded interactions spanning heavy- and light-chain V, D, and J gene segments.

### DMS Epitope Cluster 17: Coordinate binding by IGHV1-46 and IGHD2-2

IGHV1-46 was among the most strongly enriched V gene segments for Epitope Cluster 17. Analysis of the most recurrent structural interactions revealed a recurring network of interactions in 19 of 33 antibody–antigen complexes representing this epitope cluster, in which a network of germline-encoded residues from IGHV1-46 and likely-IGHD2-2 (<u>Y33</u>, <u>I50</u>, <u>N52</u>, <u>S56</u>, <u>S</u>/T<u>58</u>, <u>V</u>/I<u>98/99</u>, and <u>P99/100</u>) contact DMS-critical RBD residues E516, L518, and Y396 (**Figure 5C**).

As there were no PDB structures representing these interactions, we solved a crystal structure of the BD56-188 Fab in complex with wild-type RBD (PDB ID: 11GF) at 2.76 Å resolution, which confirmed the IGHV1-46 and IGHD2-2 recurrent interactions. Point mutations to IGHV1-46 Y33A, N52A, and S56A consistently abolished antibody binding, whereas control mutations outside of the predicted binding interface had no effect (**Figure 5C, Figure S6E**). These data indicate that germline-encoded residues in IGHV1-46 and IGHD2-2 are critical for selection and recognition of this immunodominant epitope.

Collectively, these examples show that the recognition of immunodominant epitopes often depends on coordinated germline-encoded residues distributed across multiple gene segments.

### Viral evolution is shaped by germline-encoded antibody recognition

If germline-encoded paratope-epitope interactions determine immunodominance, we reasoned that viral evolution should frequently target these sites to escape neutralization. As an initial case study, we evaluated data for several neutralizing monoclonal antibodies that received emergency use authorization from the U.S. Food and Drug Administration during the COVID-19 pandemic. Despite their initial efficacy, nearly all of these antibodies, including tixagevimab, bamlanivimab, etesevimab, bebtelovimab, and casirivimab, lost potency against emerging variants. Notably, each of these antibodies target immunodominant RBD epitopes using recurrent, germline-encoded interactions identified from our structural analysis (**Figures S7A and S7B**), explaining the sharp loss of neutralization upon mutation of the residues targeted by these recurrent interactions (e.g., L452R, E484A, F486V). These observations suggest that germline-encoded antibody features not only shape immunodominant responses but may also impose selective pressure on viral evolution. This hypothesis is supported by the mutational landscape of SARS-CoV-2 variants of concern and by corresponding antibody neutralization profiles (**Figure 6A**). For example, antibodies using recurrent interactions to engage L452 exhibit markedly reduced potency against Omicron BA.5 and BQ.1.1, which carry the L452R mutation^42^. Likewise, antibodies using recurrent interactions to contact E484 lose activity against Omicron variants (BA.1, BA.2, BA.2.75, BA.5, BQ.1.1) harboring E484A, an effect exacerbated by a proximal F486V mutation in BA.5 and BQ.1.1. Consistent with this, immunodominant epitopes corresponding to wild-type RBD DMS clusters 5 and 14 (defined by critical residues at positions 484 and 486, respectively) are diminished or absent in the BA.5 dataset, suggesting complete evasion of these antibody responses (**Figure S1B and S7C**). These patterns indicate that germline-driven antibody responses can contribute to the convergent emergence of specific RBD mutations across Omicron sublineages^18^, and that viral escape may frequently be achieved by disrupting germline-encoded binding interfaces.

**Figure 6.**
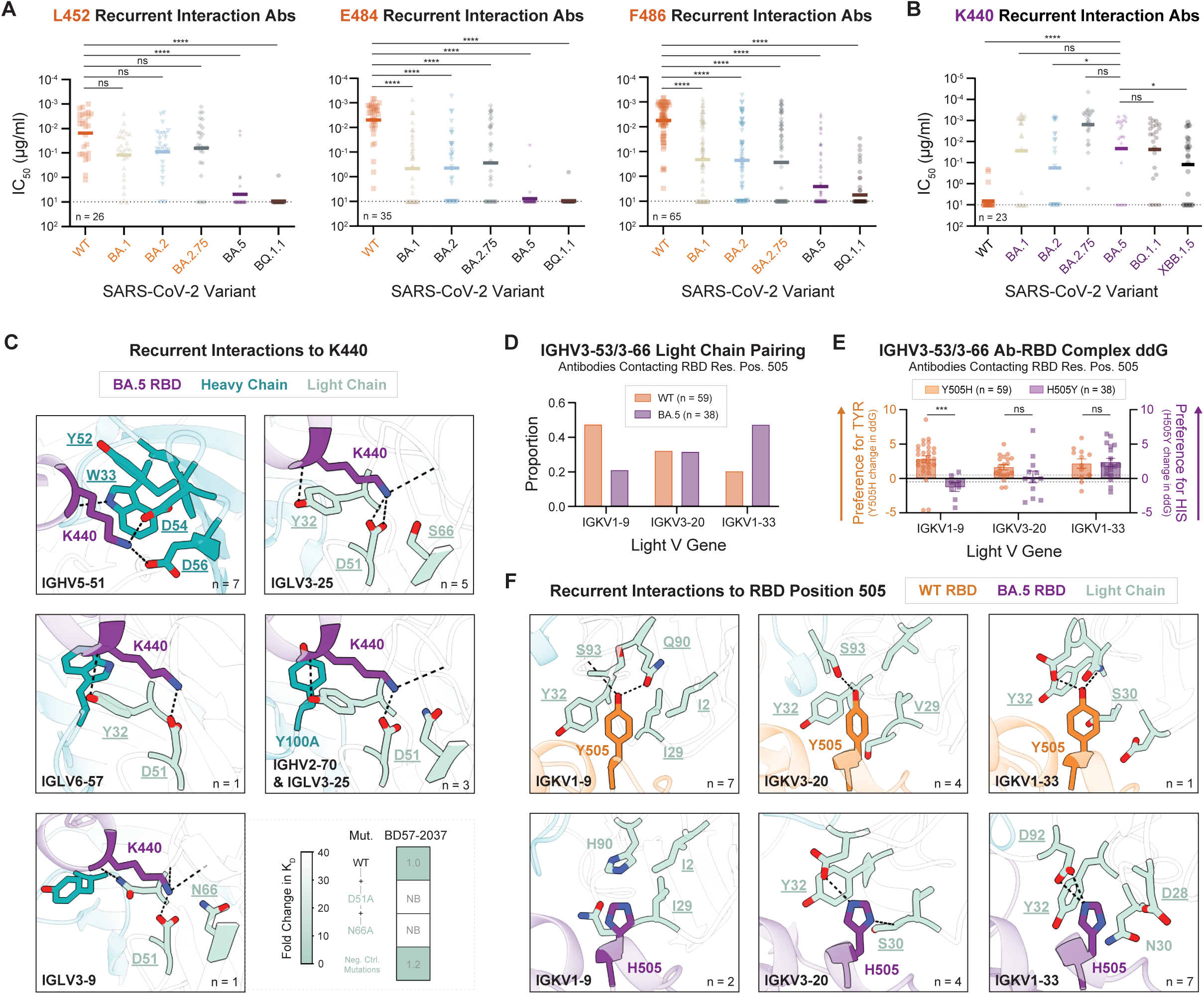
Recurrent interactions underlying the loss, formation, and maintenance of immunodominant epitopes in SARS-CoV-2 RBD. (**A**) Neutralization potency of antibodies that use recurrent interactions to target specific wild-type RBD residues across SARS-CoV-2 variants. Variants containing the specified residue are labeled in orange. Data are represented as the geometric mean IC50 and analyzed by Kruskal-Wallis with Dunn’s multiple comparisons test (****p<0.0001). (**B**) Neutralization potency of antibodies using recurrent interactions to target the BA.5 RBD N440K mutation across SARS-CoV-2 variants. SARS-CoV-2 variants containing residue K440 are labeled in purple. Data are represented as the geometric mean IC50 and analyzed using Kruskal-Wallis with Dunn’s multiple comparisons test (*p<0.05; ****<0.0001). (**C**) Recurrent interactions to the BA.5 RBD N440K mutation (purple) as observed in unique-lineage antibody-antigen complexes across BA.5 DMS epitope clusters. Dashed lines indicate the presence of hydrogen bonds or salt bridges between displayed antigen and antibody residues. Antibody residues participating in recurrent interactions are labeled in teal and pale green for the heavy chain and light chain, respectively. Fold change in binding affinity of mutant antibodies (relative to the parent antibody), measured by biolayer interferometry, is shown for a representative IGLV3-9 antibody engaging BA.5 RBD residue K440 following point mutations to the recurrent interaction (see also Figure S6). NB = No detectable binding (**D**) Proportion of IGHV3-53/3-66 antibodies pairing with highly enriched light chain V gene segments (FDR < 1x10^-7^) that recognize either wild-type RBD residue Y505 or BA.5 RBD residue H505 as a ddG hotspot (see Methods) in predicted structures. (**E**) Change in antibody-RBD complex ddG, as calculated by Rosetta, for IGHV3-53/3-66 antibodies that recognize either Y505 or H505 as a ddG hotspot following in silico Y505H or H505Y mutations. Data are presented as the mean change in the median complex ddG ± SEM following triplicate ddG calculations per antibody, and were analyzed by multiple Mann-Whitney followed by Bonferroni-Dunn multiple comparisons test (***p<0.001). (**F**) Recurrent interactions with RBD residue 505 across wild-type (top; orange) and BA.5 RBD (bottom; purple) as encoded by IGHV3-53/3-66 antibodies pairing to enriched IGKV1-9, IGKV3-20, and IGKV1-33 light chains. Dashed lines denote hydrogen bonds between displayed antigen and antibody residues. Antibody residues involved in recurrent interactions are labeled in teal and pale green for the heavy and light chains, respectively.

Conversely, viral evolution can create new immunodominant epitopes by introducing residues that are newly recognized by germline-encoded motifs. Comparative analysis of wild-type and BA.5 RBD DMS data revealed a novel BA.5 immunodominant epitope (DMS Cluster 5) centered on residue K440, created by the N440K mutation (**Figures 6B and 6C**). This epitope has no analog in wild-type RBD (**Figures 1B, S1B, and S7C**). Structural analysis identified several recurrent interactions targeting K440. Antibodies employing these motifs exhibited potent neutralization of Omicron variants but failed to neutralize wild-type SARS-CoV-2, which carries an asparagine at this position (**Figure 6B**). Many of these recurrent interactions correspond to previously described lysine-specific GRAB motifs, and their interactions share common molecular features, including analogous van der Waals contacts, hydrogen bonds, and salt bridges (**Figure 6C**). This finding demonstrates that viral mutations can generate novel immunodominant epitopes by creating binding opportunities for germline-encoded motifs.

Finally, we asked whether previously disrupted immunodominant epitopes could be rescued through alternative sets of germline-encoded interactions. A large number of antibodies featuring IGHV3-53/3-66 gene usage^43,44^ target an epitope encompassing RBD residue Y505, which is mutated to histidine (Y505H) in all Omicron subvariants. Of these antibodies, those that bound wild-type RBD predominantly used the light chain IGKV1-9, whereas those recognizing BA.5 preferentially used IGKV3-20 or IGKV1-33 (**Figure 6D**). To define the molecular basis for this shift, we analyzed antibody–antigen complex structures containing IGKV1-9, IGKV1-33, or IGKV3-20-encoded recurrent interactions targeting RBD position 505, and modeled changes in binding energy upon mutation (i.e. Y505H or H505Y) using Rosetta. We found that IGKV1-9 is optimized for tyrosine binding, whereas IGKV1-33 and IGKV3-20 can accommodate both tyrosine and histidine residues (**Figures 6E and 6F**). These results explain the observed shift in light-chain usage in BA.5-binding antibodies and illustrate how germline-encoded recognition can reconfigure to preserve binding as viral epitopes evolve. These data reveal that germline antibody constraints collectively shape how immunodominant epitopes are lost, remade, and reshaped through viral evolution.

### Limitations of the study

Several limitations should be considered. First, although the SARS-CoV-2 RBD provides an unusually rich and well-characterized dataset, it represents a single antigen. Accordingly, the extent to which these findings generalize to other antigens remains to be established. That said, there are reasons to anticipate broad applicability. Antibodies targeting the Spike subdomains NTD and S2 also exhibit strong V gene-usage biases, albeit with distinct V gene pairings that likely reflect differences in antigen structures (**Figure 1A, Figure S1A**). In addition, over 100 of the recurrent antibody-antigen interactions observed in this study are also present in non-RBD antigen contexts in the PDB, supporting the potential generality of the germline-encoded binding motifs described here. Second, the frequency of confident AF3 predictions for antibody-RBD complexes was likely influenced, in part, by the inclusion of related antibody-RBD complexes in the AF3 training set. However, the predictions did not simply reproduce binding models already seen in the PDB. For example, the recurrent interactions made by IGHV1-8 in DMS Epitope Cluster 10 and by IGHV1-46 in DMS Epitope Cluster 17 are not present in existing PDB structures. We validated these interactions experimentally by determining crystal structures and performing targeted point mutagenesis, supporting their biological relevance and the accuracy and utility of our AF3 dataset.

## Discussion

Our study provides a systematic, large-scale analysis of antibody responses to a model antigen and reveals a unifying principle: epitope selection is largely dictated by germline-encoded features of the antibody repertoire. The prevailing view in immunology has long emphasized the central role of somatic diversity, particularly within the HCDR3, in determining antigen specificity. However, by integrating antibody sequence data, deep mutational scanning, structural modeling, and experimental characterization, we provide evidence that the majority of antibodies targeting the SARS-CoV-2 RBD rely on conserved germline-encoded features, rather than diverse somatically generated ones, to recognize their epitopes. We show that germline-encoded residues from V, D, and J gene segments predominate in the most recurrent and energetically significant antibody-RBD interactions. Importantly, even when HCDR3 motifs appear to contribute to specificity, they are largely derived from germline-encoded D and J segments rather than from sequences generated during V(D)J recombination and junctional diversity. The logical conclusion is that antibody immunodominance is not solely an emergent property of antigen structure, but rather a predictable, hard-wired feature of the adaptive immune response. This conclusion aligns with prior observations that different species exposed to the same antigen tend to target distinct immunodominant epitopes^15^, likely reflecting differences in immunoglobulin gene segment repertoires.

Altogether, our findings lead us to propose that germline-encoded sequences within V, D, and J gene segments drive immunodominance in two related ways. First, for cognate epitopes these sequences provide the baseline affinity necessary for entry into the germinal center reaction. Second, because these sequences are more abundant than somatically diversified variants, they expand the pool of naïve B cell precursors available for selection - a mechanism supported by reconstruction experiments demonstrating that increasing a specific precursor frequency enhances an antigen’s ability to elicit the corresponding antibody^13^. Thus, germline sequences, by impacting both baseline affinity and precursor abundance, influence immunodominance.

We identified 104 examples of germline-encoded amino acid-binding (GRAB) motifs across 39 V gene segments, most of which are newly reported. Combined with our prior findings^15^, we therefore have data on germline-encoded binding preferences for 44 human V gene segments, approximately a quarter of the total repertoire of human V gene segments. These GRAB motifs often recognize basic amino acids, including lysine and arginine (**Figure 4A and Supplementary Data**). For a germline-encoded interaction to drive epitope selection, it must contribute sufficient binding free energy to surpass the threshold required for positive selection and clonal expansion within the germinal center. Side chains such as lysine and arginine may be particularly well suited to this role, as they offer opportunities for both electrostatic and hydrophobic interactions. In contrast, epitopes dominated by smaller or less chemically diverse residues are likely to require more extensive contact networks to achieve comparable binding energies, and thereby impose stricter geometric constraints on recognition. Beyond these 104 GRAB motifs, we identified numerous additional recurrent germline-encoded interactions that we currently observe only in the context of SARS-CoV-2 RBD. However, we anticipate that with additional future structural data on antibody-antigen complexes, the true degree of modularity of these interactions will become apparent. A more comprehensive catalog of germline configurations that preferentially recognize specific antigenic features will be essential for understanding and predicting which structural features in an antigen are likely to be immunodominant.

Our findings also provide insight into how antibody-mediated selection pressures shape viral evolution. Because antibodies recognize immunodominant epitopes through germline-encoded interactions, mutations at these epitopes can disrupt binding for large fractions of the antibody response across populations. This helps explain the repeated emergence of mutations at specific residues (e.g., L452, E484, F486) across SARS-CoV-2 variants.

Conversely, viral mutations can create new immunodominant epitopes by introducing residues that are compatible with germline-encoded motifs. The emergence of a novel epitope centered on K440 in Omicron BA.5 exemplifies how viral evolution can exploit the structure of the antibody repertoire to reshape immune responses. Together, these observations highlight a dynamic interplay between germline-encoded immune recognition and viral antigenic drift.

Because the antibody repertoire is assembled from a finite set of V, D, and J gene segments, the space of germline-encoded binding preferences is likewise finite, rendering the prediction of immunodominant epitopes a computationally tractable problem. Such predictive capacity would have important implications for the rational design of vaccines and therapeutics. Vaccine antigens could be engineered to preferentially engage or avoid specific germline features to improve vaccine efficacy. Additionally, understanding germline biases could inform germline-targeting vaccine strategies to elicit desired antibody lineages. Predicting immunodominant epitopes would also aid the design of protein therapeutics that minimize anti-drug antibody responses.

In summary, our work establishes germline-encoded binding as a dominant force shaping the majority of the antibody response and resulting in the phenomenon of immunodominance. These findings redefine the balance between germline- and non-germline-encoded contributions to antibody specificity and suggest that the architecture of the germline repertoire imposes fundamental constraints on immune recognition. Future studies extending this approach to diverse antigens will be critical to fully decode the molecular rules of immunogenicity and harness this information to guide vaccine and therapeutic design.

## Resource availability

All data necessary to interpret our findings, including the Supplementary Data, are available on the Harvard Dataverse https://doi.org/10.7910/DVN/U99KRL (Unpublished Dataset Preview URL: https://dataverse.harvard.edu/previewurl.xhtml?token=7ded3d60-7926-4a18-a986-9cf086bd503a). All reasonable requests for materials will be fulfilled. Correspondence and material requests should be addressed to.

## Acknowledgments

We thank Frederick Alt, Han Altae-Tran, Nathaniel Bennett, Yunlong Cao, Nir Hacohen, David Juergens, Indrek Kalvet, Elijah Mena, Joseph Watson, and Duane Wesemann for helpful discussions; Luki Goldschmidt and Patrick Vecchiatto for maintenance of and assistance with the computing cluster at the University of Washington Institute for Protein Design; and Jeremy Yuenger for assistance with the Harvard Dataverse. We thank the Resource for Biocomputing, Visualization, and Informatics at the University of California, San Francisco, with support from NIH P41-GM103311 for help with molecular graphics. We thank Thomas W. Linsky for help with Rosetta software. This manuscript was reviewed and released by Lawrence Livermore National Laboratory (LLNL) (LLNL-JRNL-2011517-DRAFT).

## Author contributions

E.L.S. and S.J.E. conceived the study. E.L.S. prepared DMS datasets and analyzed gene segment usage with assistance from I.-H.L. Z.Y. developed SWIM algorithm; Z.Y. and E.L.S. identified HCDR3 sequence motifs. M.H.W., E.L.S., and M.B.G. generated AF3-predicted structures. M.H.W. performed structural clustering of antibody-RBD complexes. E.L.S. identified recurrent structural interactions with assistance from E.S. E.S. identified Rosetta ddG hotspots with assistance from B.C. and analyzed evolution of immunodominant epitopes from wild-type to BA.5 SARS-CoV-2. A.D. measured changes in binding affinity upon mutation of recurrent structural interactions and obtained crystal structures with guidance from D.I and supervision from J.-P.J. S.J.E. and D.B. supervised the study. E.L.S., E.S., and S.J.E. prepared the manuscript with contributions from M.H.W. and Z.Y. S.J.E. and D.B. reviewed and edited the manuscript with input from all co-authors.

## Declaration of generative AI and AI-assisted technologies

Throughout this study, large language models including ChatGPT, Google Gemini, and Claude were used in developing code and editing the manuscript.

## Supplemental information titles and legends

### Supplementary Discussion

#### Point #1

Antibodies in different DMS epitope clusters exhibit varying patterns of neutralization potency (**Figure S7D**).

#### Point #2

Many of the HCDR3 motif clusters also featured biased V gene segment usage (**Supplementary Data**).

Upon generation of the AF3 predicted dataset, we examined whether residues within conserved HCDR3 motifs generally contacted DMS-critical residues or ddG hotspots on the RBD, which could account for their conservation. In almost all cases, they did (**Supplementary Data**).

#### Point #3

We note that none of IgLV3-10, IgLV3-25, IgLV6-57, IgLV3-1, or IgLV5-37, i.e. genes that harbor lysine-specific GRAB motifs similar to that in IgLV3-21, share the <u>S</u>/N<u>32</u>, <u>Y49</u>, <u>D/Y50</u>, and <u>D53</u> residues predicted to contact RBD R357. In particular, IgLV3-10, IgLV3-25, and IgLV3-1 have positively charged residues in one of these positions that could repel RBD R357. (Germline-encoded residues at these positions are as follows: IgLV3-10: <u>Y32</u>, <u>Y49</u>, <u>E50</u>, <u>K53</u>; IgLV3-25: <u>Y32</u>, <u>Y49</u>, <u>K50</u>, <u>E53</u>; IgLV3-1: <u>Y32</u>, <u>Y49</u>, <u>Q50</u>, <u>K53</u>; IgLV6-57: <u>Y32</u>, <u>Y49</u>, <u>E50</u>, <u>Q53</u>; IgLV5-37: <u>N32</u>, <u>Y49</u>, <u>Y50</u>, and <u>G53</u>^34–36.)^

#### Point #4

One specific residue, N32 in IgLV3-21, was not annotated as germline-encoded. However, since the S-to-N substitution requires multiple nucleotide changes and is therefore less likely to arise through somatic hypermutation, it is plausible that this N32 is germline-encoded in an as-yet-undescribed IgLV3-21 allele, potentially enriched in the Chinese population from which the dataset was derived.

## Supplementary Figure Legends

**Figure S1. Antibodies to immunodominant wild-type and BA.5 RBD epitopes exhibit biased V gene segment usage, suggesting germline-encoded recognition.** (**A**) Heavy-light V gene segment usage of antibodies that recognize the wild-type RBD, NTD, and S2 subdomains of SARS-CoV-2 spike, as in Figure 1A, with labels. (**B**) BA.5 RBD DMS dataset, clustered and visualized as in Figure 1B. (**C and D**) Heavy-light V gene segment usage of antibodies in each (C) wild-type RBD or (D) BA.5 DMS Epitope Cluster, as in Figure 1C, with labels.

**Figure S2. There is a strong association between gene segment usage and epitope specificity.** (**A and B**) Comparison of the epitopes of antibodies with highly enriched heavy-light V gene segment pairs. To visualize and cluster antibodies based on their wild-type RBD DMS epitope maps, we applied t-distributed stochastic neighbor embedding (t-SNE) to reduce the feature space to two dimensions, then performed k-means clustering. This clustering method is the same as was used for Figure 1B. Points representing antibodies with particular overrepresented V gene segment pairs are shown in color, while points representing all other antibodies are in light grey. (**C**) Average wild-type RBD DMS data for unique-lineage antibodies that share overrepresented heavy-light V gene segment pairs. A DMS escape score of ≥ 0.3 was applied as a threshold to define critical RBD epitope residues for each antibody (see Methods). Then the DMS data, now converted to 0 or 1 binary values, were averaged for all antibodies with the indicated heavy-light V gene segment pairs. To limit figure size, only heavy-light V gene segment pairs enriched over their usage in the naïve antibody repertoire with FDR < 1.0 x 10^-7^ were plotted. Rows in the heatmap were clustered using hierarchical clustering. Rows representing antibodies with overrepresented V gene segment pairs mentioned in (A) and (B) are shown in color and labeled with grey circled numbers at left that correspond with grey circled numbers in the top left corner of plots in (A).

**Figure S3. Unsupervised HCDR3 clustering reveals conserved sequence motifs largely germline-encoded by D and J gene segments.** (**A**) UMAP representations illustrate antibody sub-clusters reflecting conserved HCDR3 sequence motifs. (**B**) HCDR3 motifs identified within wild-type RBD DMS epitope clusters by SWIM analysis.

**Figure S4. Structural clustering of antibody–RBD complexes recapitulates DMS epitope clusters. (A)** All-by-all clustering of AF3-predicted structural models of antibody-wild-type RBD complexes and PDB structures of antibody-antigen complexes where the antibody is annotated in the CoV-AbDab to bind to any pre-Omicron SARS-CoV-2 RBD. Shown from left to right are the RMSD matrix from all-by-all structural alignments, the dataset of origin for each antibody, the assigned structural cluster, and DMS Epitope Cluster (defined as in Figure 1B and Supplementary Data). The order of antibodies on the x- and y- axes are identical across all subpanels. Because 22 DMS epitope clusters were defined, 22 structural clusters were specified. **(B)** Number of antibody-antigen complexes from (A) corresponding to each DMS Epitope Cluster that fall within each wild-type RBD Structural Cluster. **(C)** As in (A) but using AF3-predicted structural models of antibody-BA.5 RBD complexes and PDB structures of antibody-antigen complexes where the antibody is annotated in the CoV-AbDab as binding to any Omicron SARS-CoV-2 RBD. **(D)** As in (B) but using the antibody-antigen complexes from (C).

**Figure S5. Analysis of critical epitope residues identified by deep mutational scanning and structural modeling.** (**A**) Number of DMS-critical residues per antibody identified. (**B**) Number of DMS-critical residues and ddG hotspots identified in the interface of each antibody-antigen complex structural model. (**C**) Overlap between DMS-critical residues and ddG hotspots for each DMS Epitope Cluster.

**Figure S6. Features of recurrent antibody-antigen interface motifs.** (**A**) Composition of paratope and epitope residues at antibody-antigen interfaces, within recurrent interactions, and in GRAB motifs. (**B-F**) Effects of recurrent-interaction residue mutations on Fab binding affinity, as measured by biolayer interferometry. Each row represents binding affinity data for one Fab and point mutants thereof. The recurrent interaction predicted to be present in the given antibody-RBD complex is shown at right. Negative control mutations target residues predicted to lie outside of the antibody-antigen interface. Orange-colored binding curves are for Fabs to wild-type RBD, whereas purple-colored binding curves are for Fabs to BA.5 RBD.

**Figure S7. Loss, formation, and maintenance of immunodominant epitopes in SARS-CoV-2 RBD.** (**A**) Gene usages of therapeutic monoclonal antibodies that use recurrent interactions are highlighted in plot from Figure 1A. (**B**) Recurrent interactions made by therapeutic monoclonal antibodies. (**C**) Extent of DMS epitope cluster overlap between wild-type and BA.5 RBD, as defined by the Jaccard index between their respective sets of DMS-critical residues. (**D**) Neutralization activity of antibodies across wild-type and BA.5 RBD DMS Epitope Clusters.

## Methods

### Curation of Unique-lineage Antibody Sequences from the CoV-AbDab and DMS Datasets

We downloaded the CoV-AbDab (version 13 June 2023) and filtered it to only include human antibodies with unique heavy chain variable domain sequences and non-null light chain variable domain sequences. To exclude potentially clonally related antibodies in order to achieve unbiased detection of enriched gene segments, antibodies were first grouped by heavy and light V and J gene segment usage. Any antibody with unique gene segment usage was retained. Then, within each gene segment usage group with more than one member, we used KA-Search^45^ to calculate all-by-all HCDR3 identities.

Any antibody with HCDR3 identity < 0.7 to all other members of the group was retained. Of the antibodies with ≥ 0.7 HCDR3 identity to at least one other antibody in the same group, if the HCDR3 length or source was different from those of all the matches, we retained it. Of the remaining antibodies with ≥ 0.7 HCDR3 identity, identical HCDR3 length, and identical source to at least one another antibody in the same group, we retained one example of each group. This yielded “unique-lineage” antibodies. We performed the same filtering steps for the antibodies from the BA.5 RBD DMS dataset, as these were not included in the version of the CoV-AbDab we used.

### Calculation of V Gene Segment Usage

From the set of unique-lineage CoV-AbDab antibodies, we identified ones that bind to the wild-type RBD by filtering for rows with the value “S; RBD” in the field “Protein + Epitope” and that included the value “SARS-CoV2_WT” in the field “Binds to”. We identified antibodies to the NTD by filtering for rows with the values “S; NTD” or “S: NTD” in the field “Protein + Epitope”. We identified antibodies to the S2 domain by filtering for rows with the values “S; S2” or “S; S2 Fusion Peptide” or “S; S2 Stem Helix” or “S; Stem Helix” in the “Protein + Epitope” field.

To establish a baseline for V gene segment usage, we used a dataset of 661,780 antibody sequences from the Observed Antibody Space database annotated as originating from naïve human B cells. We used a one-sided Fisher’s exact test with Benjamini-Hochberg False Discovery Rate (FDR) correction, comparing the counts of antibodies with each given V gene segment or paired heavy-light V gene segments in the set of interest compared to in the set of naïve antibodies. V gene segment(s) with FDR < 0.05 were considered to be significantly enriched in the set of interest relative to in the naïve human antibody repertoire.

### DMS Data Processing and Clustering

DMS escape scores and neutralization IC50s against SARS-CoV-2 variants were obtained from https://github.com/jbloomlab/SARS2-RBD-escape-calc (version January 2024). Unique-lineage antibodies with DMS data (i.e., antibodies present in both the unique-lineage CoV-AbDab dataset and the wild-type RBD DMS dataset, or unique-lineage antibodies from the BA.5 DMS dataset) were used for all subsequent analysis. To visualize and cluster antibodies based on their wild-type RBD or, separately, BA.5 RBD DMS epitope maps, we first applied t-distributed stochastic neighbor embedding (t-SNE) to reduce the feature space to two dimensions. The resulting two-dimensional embeddings were then subjected to k-means clustering to identify distinct groups of antibodies. We specified 22 clusters to approximately reflect the observed diversity of antibody responses.

### Calculation of Average DMS Epitopes

Wild-type RBD DMS data for each antibody were processed by applying a threshold of ≥ 0.3 to define critical epitope residues, resulting in binary (0 or 1) values for each position along the RBD. Binarized data were averaged across all unique-lineage antibodies with a given heavy-light V gene segment pair to obtain the proportion of antibodies that rely on each residue of the RBD for binding. For the BA.5 RBD DMS data, a threshold of ≥ 0.5 was used to define critical RBD epitope residues.

### HCDR3 Sequence Motif Analysis

The SWIM algorithm starts by assigning a distance between every pair of amino acids (AAs), AA_i_ and AA_j_ (i,j = 1,2,…,20). This distance, d(AA_i_,AA_j_), is derived from the BLOSUM62 matrix using the following formula:

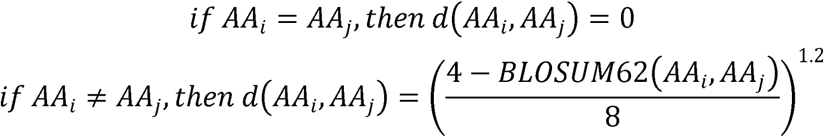

*Where the number 8 in the denominator normalizes d(AAi,AAj) to between 0 and 1, and raising the normalized distance to the power of 1.2 emphasizes similarities between AAs to allow better HCDR3 clustering*.

The algorithm uses this AA-level distance to compute a distance between any given pair of HCDR3s. First, each IMGT-defined HCDR3 is trimmed to remove the first two and the last two AAs (these four amino acids are well conserved across all HCDR3s as position 0 and its length denoted as *L*, the algorithm then locates the centermost 4- AA window starting at position 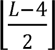. From this central position, the window is slid to the right one AA at a time for up to 4 times, or until the end of the HCDR3 is reached. Similarly, the window is slid to the left for up to 4 times. This procedure extracts up to 9 4-AA windows from each HCDR3. For any pair of HCDR3s, the algorithm considers distances between all pairs of windows (*W*_1_ and *W*_2_) extracted from the two HCDR3s, with the window-to-window distance defined as:

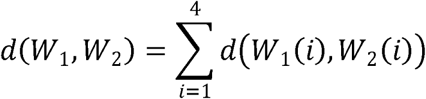

Where *w*_1_(i) denotes the ith *AA* in *w*_1_. Finally, the algorithm defines the distance between the two HCDR3s as:

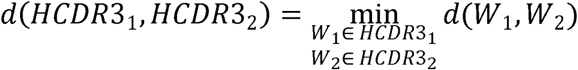

The pairwise distances between all HCDR3s being analyzed are then passed on to the UMAP dimensionality-reduction algorithm (with number of neighbors set to 15, minimum distance set to 0.1, and random state set to 42) to obtain a two-dimensional embedding that can be used to visualize sequence similarities between HCDR3s. HCDR3 clusters are then obtained in an unsupervised manner by feeding the UMAP embedding to the HDBSCAN algorithm (with minimum cluster size set to 10, minimum samples per cluster set to 10, and cluster selection epsilon set to 0).

### AlphaFold 3 Prediction of Antibody-Antigen Complexes

For all unique-lineage antibodies in the wild-type RBD and BA.5 RBD DMS datasets, we attempted AF3 predictions with the heavy and light variable regions in complex with the respective RBD antigen. The wild-type RBD sequence used for the predictions was QCVNLTTRTQLPPAYTNSFTRGVYYPDKVFRSSVLHSTQDLFLPFFSNVTWFHAIHVSGTNGTKRFDNPVLPFNDGVYFASTEKSNIIRGWIFGTTLDSKTQSLLIVNNATNVVIKVCEFQFCNDPFLGVYYHKNNKSWMESEFRVYSSANNCTFEYVSQPFLMDLEGKQGNFKNLREFVFKNIDGYFKIYSKHTPINLVRDLPQGFSALEPLVDLPIGINITRFQTLLALHRSYLTPGDSSSGWTAGAAAYYVGYLQPRTFLLKYNENGTITDAVDCALDPLSETKCTL. The BA.5 RBD sequence used was RVQPTESIVRFPNITNLCPFDEVFNATRFASVYAWNRKRISNCVADYSVLYNFAPFFAFKCYGVSPTKLNDLCFTNVYADSFVIRGNEVSQIAPGQTGNIADYNYKLPDDFTGCVIAWNSNKLDSKVGGNYNYRYRLFRKSNLKPFERDISTEIYQAGNKPCNGVAGVNCYFPLQSYGFRPTYGVGHQPYRVVVLSFELLHAPATVCGPKKSTNLVKNKCVNF. Antibody- antigen complexes were predicted using one seed. For each antibody-antigen complex, the output structural model with the highest ipTM was collected. Then we filtered for high-confidence predictions by selecting structural models with ipTM values ≥ 0.8. Structural models were renumbered using the Chothia numbering system for the antibody chains and the appropriate residue positions within the RBD.

### Curation of Antibody-Spike Structures from the PDB

Experimentally determined structures of antibodies in the CoV-AbDab were identified from the CoV-AbDab summary file, and Chothia-numbered PDB files of these structures were retrieved from the SAbDab. Annotations from the CoV-AbDab and SAbDab were merged to collect all relevant information. The dataset was filtered for entries containing heavy, light, and target components and with the value “S; RBD” in the field “Protein + Epitope”. Obvious non-human entries with the values “macaca mulatta” or “oryctolagus cuniculus” in the field “heavy_species” were excluded. If multiple PDB structures were present for a given antibody, only the highest resolution structure was selected. All antibody structures retrieved from the SAbDAb were standardized to contain two chains prior to any downstream analyses, including single-chain variable fragment structures, which were pre-processed to separate heavy and light chain components according to SAbDAb-annotated chain labels.

### All-by-all Structural Clustering of Antibody-Antigen Complexes

For each pair of antibody-antigen complexes, the antigens were first aligned based on sequence, then structurally aligned on shared residue positions using the Kabsch algorithm. For PDB structures involving multiple instances of the same antibody-RBD complex (for instance, a structure involving a spike trimer with three identical antibody-RBD complexes), only one instance, selected with no particular order, was used. As AF3 predictions of the RBD resulted in structural models in which the ends were predicted as unstructured regions with very low confidence, only the middle 75% of the sequence was considered during alignment. To calculate RMSDs between antibody CDRs of variable lengths, we calculated RMSDs over N-Cα-C atoms of Chothia-defined residue positions shared between the given pair of antibody-antigen complexes. Hierarchical clustering was subsequently performed on CDR RMSD distances to generate 22 structural clusters.

### Rosetta-based Identification of ddG Hotspots and Analysis of DMS-ddG Overlap

AF3-predicted models of antibody-antigen complexes were analyzed computationally by Rosetta to identify key epitope-paratope interactions. Complexes were first refined using a Rosetta FastRelax protocol with coordinate constraints prior to per-residue ddG calculation. For each residue at the interface, Rosetta energy contributions of each amino acid side chain were evaluated by the beta_nov16 score function and subsequently filtered for ddG scores ≤ -1.0 to identify ddG hotspot residues.

To evaluate the quality of the AF3-predicted structural models, we calculated the percentage of DMS-critical residues found among the set of computationally identified ddG hotspots, with high percentage overlap indicative of good agreement between experimental DMS data and the predicted binding interface. As a negative control, 250 wild-type and 250 BA.5 antibodies were randomly selected from the AF3-predicted dataset (negative control antibody) and paired with a random antibody from a different epitope cluster. For each pairing, we calculated percent overlap between the DMS-critical residues of the random antibody and the ddG hotspots of the negative control antibody. Antibodies for which there were structures deposited in the PDB on or prior to the AF3 training cutoff date of September 30, 2021 were omitted from this analysis (14 antibodies to wild-type RBD excluded).

### *In silico* analysis of Y505H and H505Y mutations

IGHV3-53/3-66 antibodies recognizing either wild-type RBD residue Y505 or BA.5 RBD residue H505 as a ddG hotspot were subject to *in silico* mutagenesis followed by Rosetta ddG calculations. For each antibody-antigen complex, reciprocal Y505H (wild-type RBD) and H505Y (BA.5 RBD) mutations were introduced on relaxed structures using Rosetta, followed by scoring of the complex. Mock Y505Y and H505H mutations and ddG scoring were performed in parallel to obtain baseline binding energies for each antibody-antigen structure. All experiments were run in triplicate and median values were used to calculate the average change in complex ddG, which is defined here as the difference in complex ddG after mock and reciprocal mutation.

### Identification of Recurrent Interactions

Contacts between antibody chains and antigen were identified using the findclash command in UCSF Chimera, which uses van der Waals radii overlap to identify atom-atom contacts. We applied an overlap cutoff of -1.0 Å and hbondAllowance of 0 to identify atom-atom contacts. Atom-atom contact results were summarized to residue-residue contacts. For downstream analysis, only antibody-antigen contacts involving atoms from antibody side chains were considered.

All confident antibody-wild-type RBD AF3 predictions and antibody-RBD structures from the PDB with associated DMS data, and, separately, all confident antibody-BA.5 RBD AF3 predictions were analyzed for recurrent interactions. We first identified cases where at least two antibody residues at Chothia-defined positions interacted through their side chains with a given DMS-critical or ddG hotspot residue in the antigen in at least three unique-lineage antibodies. To reduce redundancy, if there were two recurrent interactions involving the same antigen residue, and one (the “superset”) included all the antibody residues of the other (the “subset”) and was observed in at least half of the antibodies that the subset recurrent interaction was seen in, the subset recurrent interaction was removed. Recurrent interactions were subsequently separated into further groups by the V gene segment(s) of the participating heavy and/or light chain(s).

### Identification of GRAB Motifs

All human antibody-antigen complexes in the SAbDab (version 18 Oct 2024) were analyzed for antibody-antigen contacts using the findclash command from UCSF Chimera as described above. For every given recurrent interaction observed in the RBD datasets, we checked to see if the same antibody interaction, involving the same heavy and/or light V gene segment and antibody residues (at Chothia-defined positions) was made with the same amino acid in a different antigen context in the PDB. Then, to quantify the diversity of antigen sequences represented in the PDB with the given recurrent interaction, we implemented a two-tiered sequence comparison and clustering strategy. First, pairwise antigen sequence similarity was initially assessed using a k-mer-based approach. Specifically, sequences were tokenized into overlapping k-mers (window size = 5), and cosine similarity was computed using the dot product of the resulting k-mer frequency vectors. This fast approximation filtered sequence pairs for which the similarity was below a specified threshold (85% identity). Filtered pairs were then refined using global pairwise alignment via the Needleman-Wunsch algorithm (pairwise2.align.globalxx) from the Biopython library. Sequence identity was calculated as the proportion of identical residues over the length of the shorter sequence. For each group, sequences with no other sequences exceeding the identity threshold were marked as unique. To further differentiate non-RBD-like antigens, the sequence of SARS-CoV-2 wild-type RBD was appended to each group, and pairwise similarity between each antigen sequence and the RBD was recalculated. Antigen sequences with <85% identity to the RBD and no similar counterparts in the group were counted as unique, non-RBD antigens.

### Expression and Purification of Fabs for Binding Panel

The heavy and light chain plasmids of various His-tagged Fabs were co-transfected at a 2:1 ratio in HEK293F cells grown in suspension using GIBCO FreeStyle 293 Expression Medium. The cultures were transfected using PEI-MAX (Kyfora Bio) at a cell density of 0.8 million cells per mL. Cultures were maintained at 37°C, 8% CO_2_, 70% humidity and rotating at 130 RPM. After 7 days, the supernatants of each culture were isolated through centrifugation of the cultures at 4000 x g for 15 minutes. The supernatants were then filtered through a 0.22 μm membrane and the Fabs were purified using a HisTrap Excel column (Cytiva) into a final buffer of 20 mM Tris and 150 mM sodium chloride at pH 8.0.

### Biolayer Interferometry

Biolayer interferometry experiments were conducted at 22°C using an OctetRED96 instrument (Sartorius FortéBio) to determine the binding kinetics of wild-type RBD and BA.5 RBD to various Fabs. Both the Fabs and RBDs were diluted into kinetics buffer (PBS pH 7.4, 0.001% (w/v) BSA, and 0.002% (v/v) Tween-20). Fabs at 20 μg/mL were immobilized onto Fab2G biosensors (Sartorius FortéBio) and were loaded until a BLI signal response of 1.0 nm was reached, followed by a 60 s baseline step in kinetics buffer. Biosensors were then dipped into wells containing a dilution series of either wild-type RBD or BA.5 RBD from 125 nM to 7.8 nM, and association was followed for 180 s. Biosensors were then dipped into kinetics buffer for 300 s to allow for dissociation of the RBDs from the Fabs. To analyze the kinetics data, the Octet Data Analysis software version 9.0.0.15 (Sartorius FortéBio) and GraphPad Prism 9 were used.

### Crystallization and Structure Determination

Purified wild-type RBD and anti-kappa V_H_H domain^46^ were complexed with Fabs in a 1.4:1.4:1 molar ratio, and excess wild-type RBD and anti-kappa V_H_H domain were separated from the Fab-antigen complexes using size-exclusion chromatography (Superdex 200 Increase 10/300 GL, Cytiva) in 20 mM TRIS and 150 mM sodium chloride at pH 8.

The wild-type RBD, BD56-188 Fab, and anti-kappa V_H_H domain complex was concentrated to 15 mg/mL and mixed in a 1:1 ratio with a crystallization buffer containing 0.2 M lithium sulfate, 0.1 M sodium acetate, and 30% (w/v) polyethylene glycol 8000 at pH 4.5. The resulting crystals grew at 22°C from a sitting-drop, vapor diffusion scheme. Before being cooled in liquid nitrogen, crystals were cryoprotected in crystallization buffer mixed with ethylene glycol, with a final concentration of 15% ethylene glycol.

The wild-type RBD, BD55-6641 Fab, and anti-kappa V_H_H domain complex was concentrated to 17.5 mg/mL and mixed in a 1:1 ratio of protein to crystallization buffer containing 0.5 M potassium thiocyanate and 0.1 M sodium acetate at a pH of 4.6. Sitting drops were set in a vapor diffusion scheme, and crystals grew at 22°C. Crystals were then cryoprotected in crystallization buffer mixed with ethylene glycol to a final concentration of 15% ethylene glycol before being cooled in liquid nitrogen.

Data for the wild-type RBD, BD56-188 Fab, and anti-kappa V_H_H domain co-complex crystal were collected at the 23-ID-B beamline Argonne National Laboratory Advanced Photon Source at a wavelength of 1.03320 Å and temperature of 100 K. The resulting dataset was processed using XDS (v20251103) and anisotropic diffraction was noted with diffraction limits of the principal axes being 5.21 Å, 2.69 Å, and 2.74 Å. STARANISO (v3.402) was used for anisotropic truncation and scaling with a local diffraction cut-off criterion of <I/sd(I)> set to 2.35. For the wild-type RBD, BD55-6641 Fab, and anti-kappa V_H_H domain co-complex crystal, data was collected at the 17-ID-1 beamline at the Brookhaven National Laboratory National Synchrotron Light Source II at a wavelength of 0.91990 Å and temperature of 100 K. The resulting dataset was processed and scaled using XDS (v20251103) and Aimless. The initial structures of both complexes were determined by molecular replacement using Phaser. Structural refinements were performed using PHENIX (v1.21.2-5419). The resulting models were checked and manually improved using Coot (v0.9.8.96). Final Ramachandran statistics of the wild-type RBD, BD56-188 Fab, and anti-kappa V_H_H domain co-complex structure showed that 96.5% of the residues were favored and 3.5% were allowed with 0% outliers. Final Ramachandran statistics of the wild-type RBD, BD55-6641 Fab, and anti-kappa V_H_H domain co-complex structure showed that 98.0% of the residues were favoured and 2.0% were allowed with 0% outliers. All software was accessed through SBGrid (v2.12.3).

