## Supplemental Figures for "Systems-Scale Structural Modeling Reveals the Germline Architecture of Immunodominance"

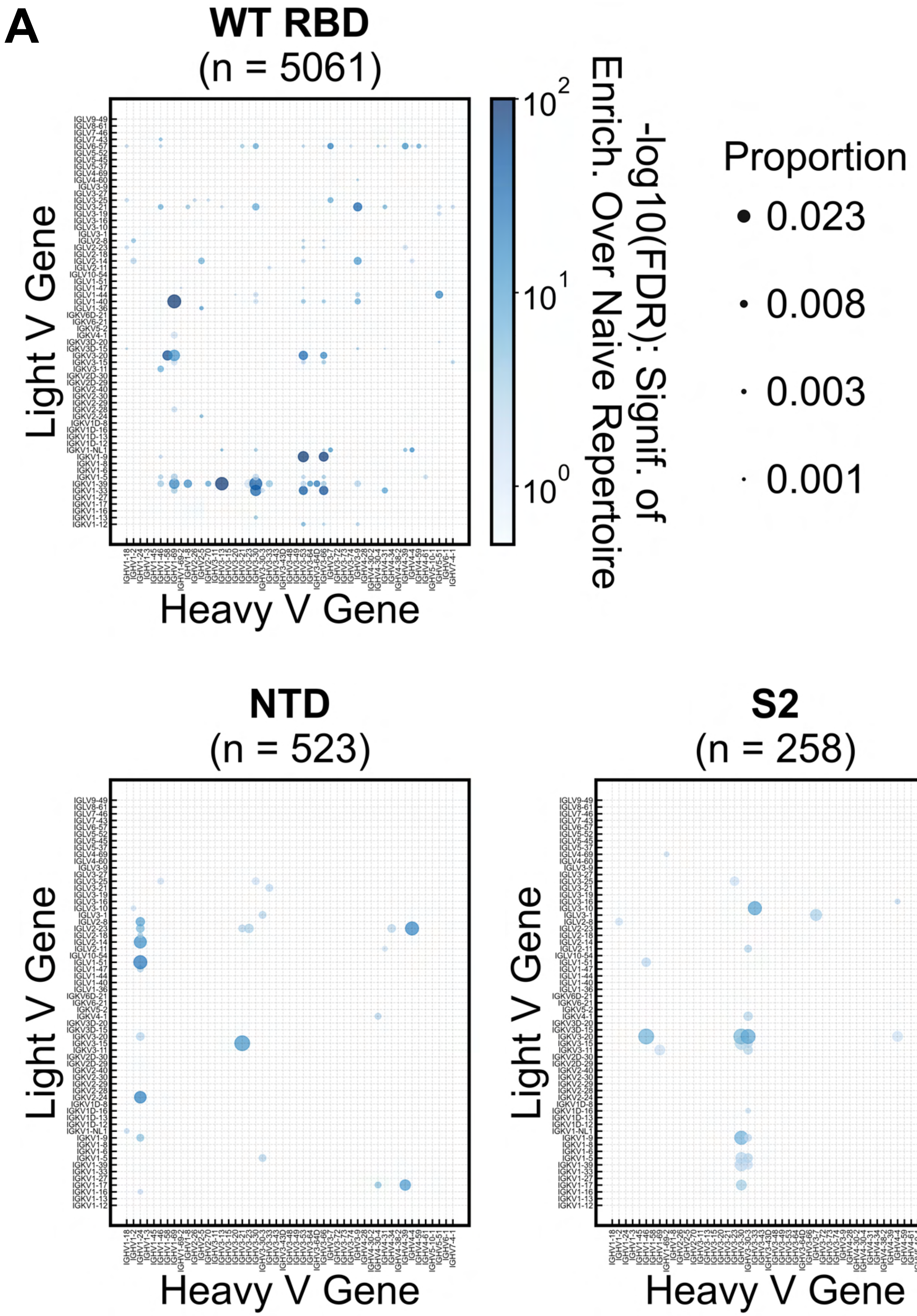
A
WT RBD
(n = 5061)
NTD
(n = 523)
S2
(n = 258)
Light V Gene
Heavy V Gene
-log10(FDR): Signif. of
Enrich. Over Naive Repertoire
$10^2$
$10^1$
$10^0$
Proportion
0.023
0.008
0.003
0.001

Figure S1

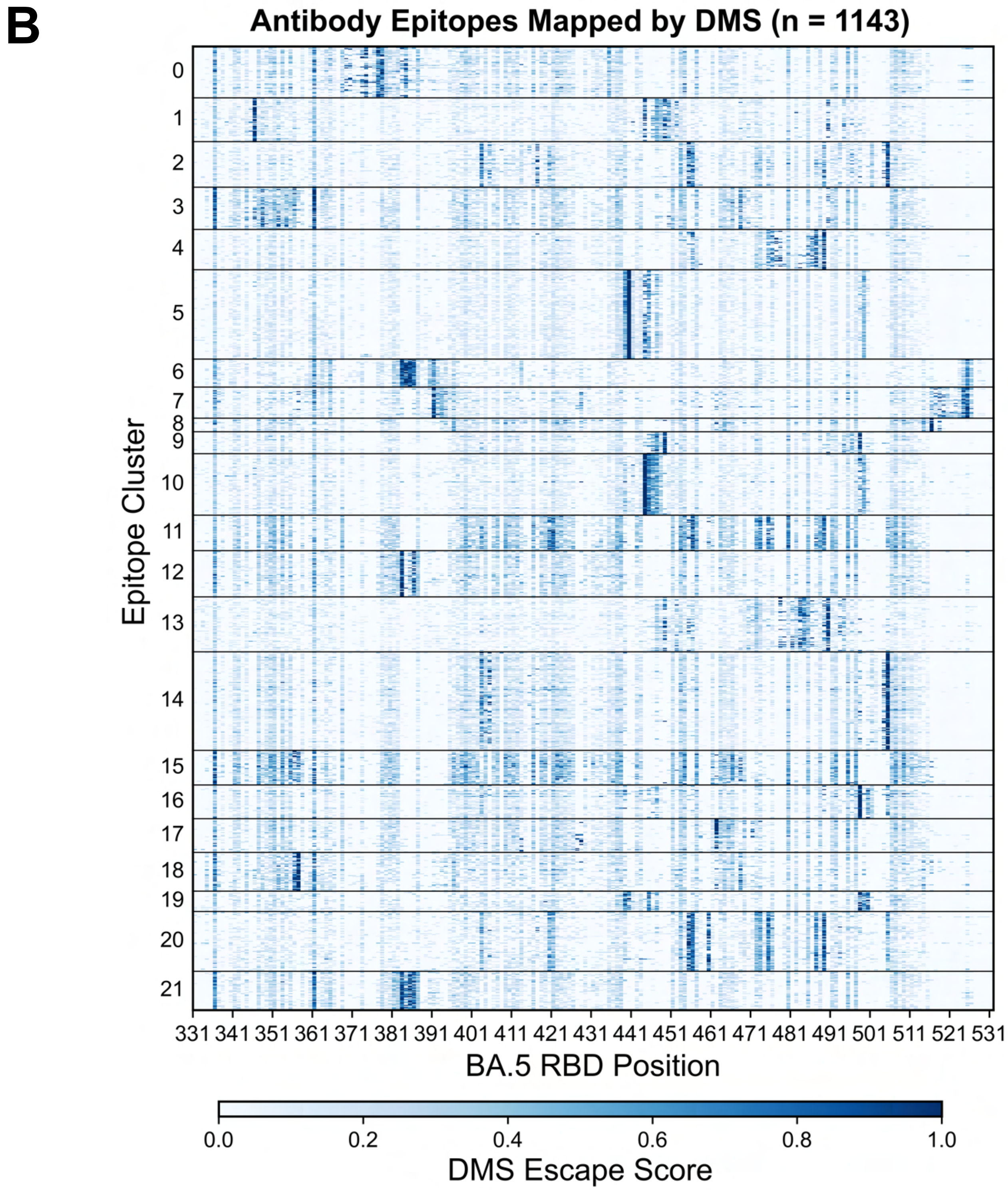


Figure S1 continued

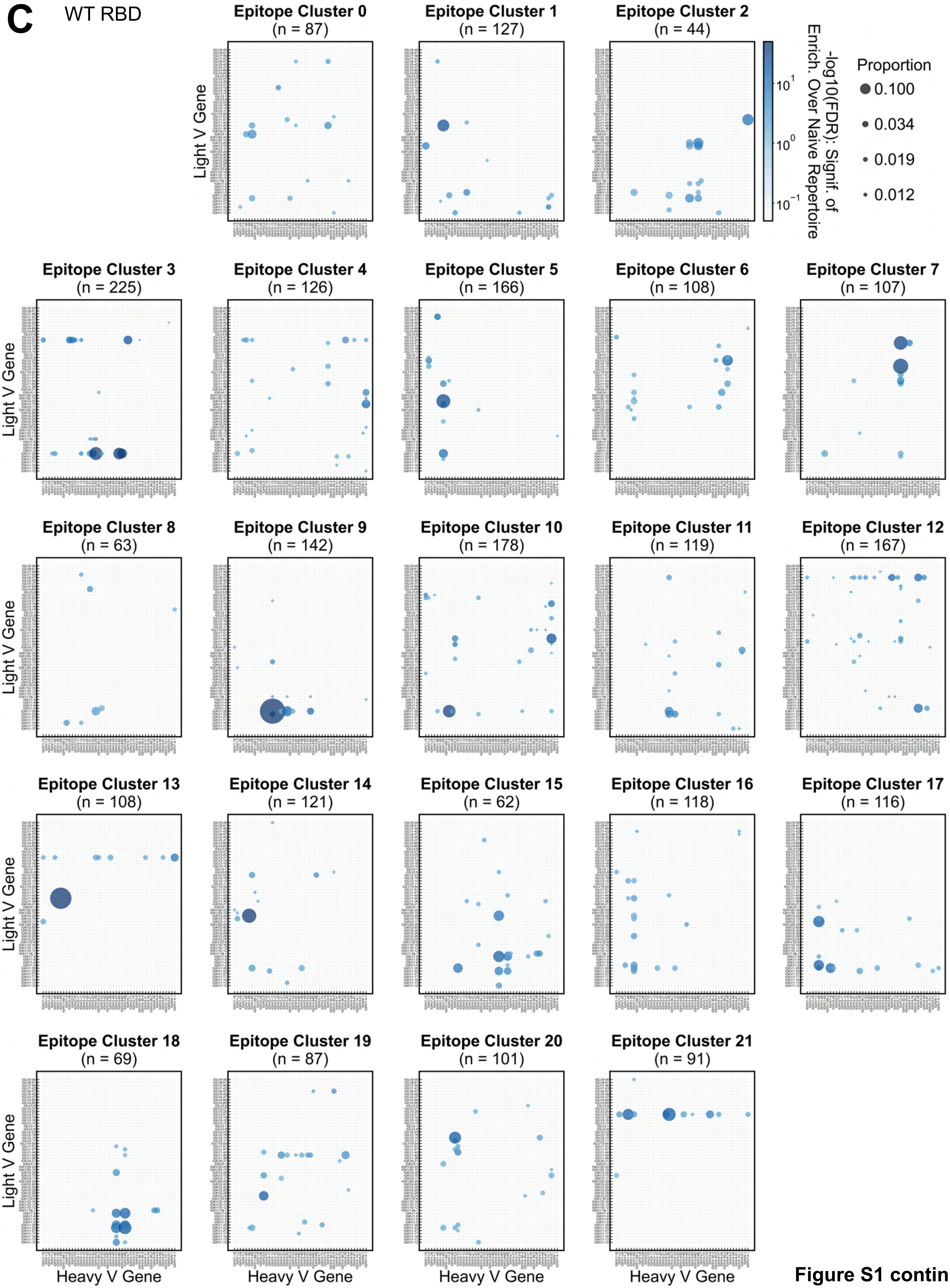


**Figure S1 continued**

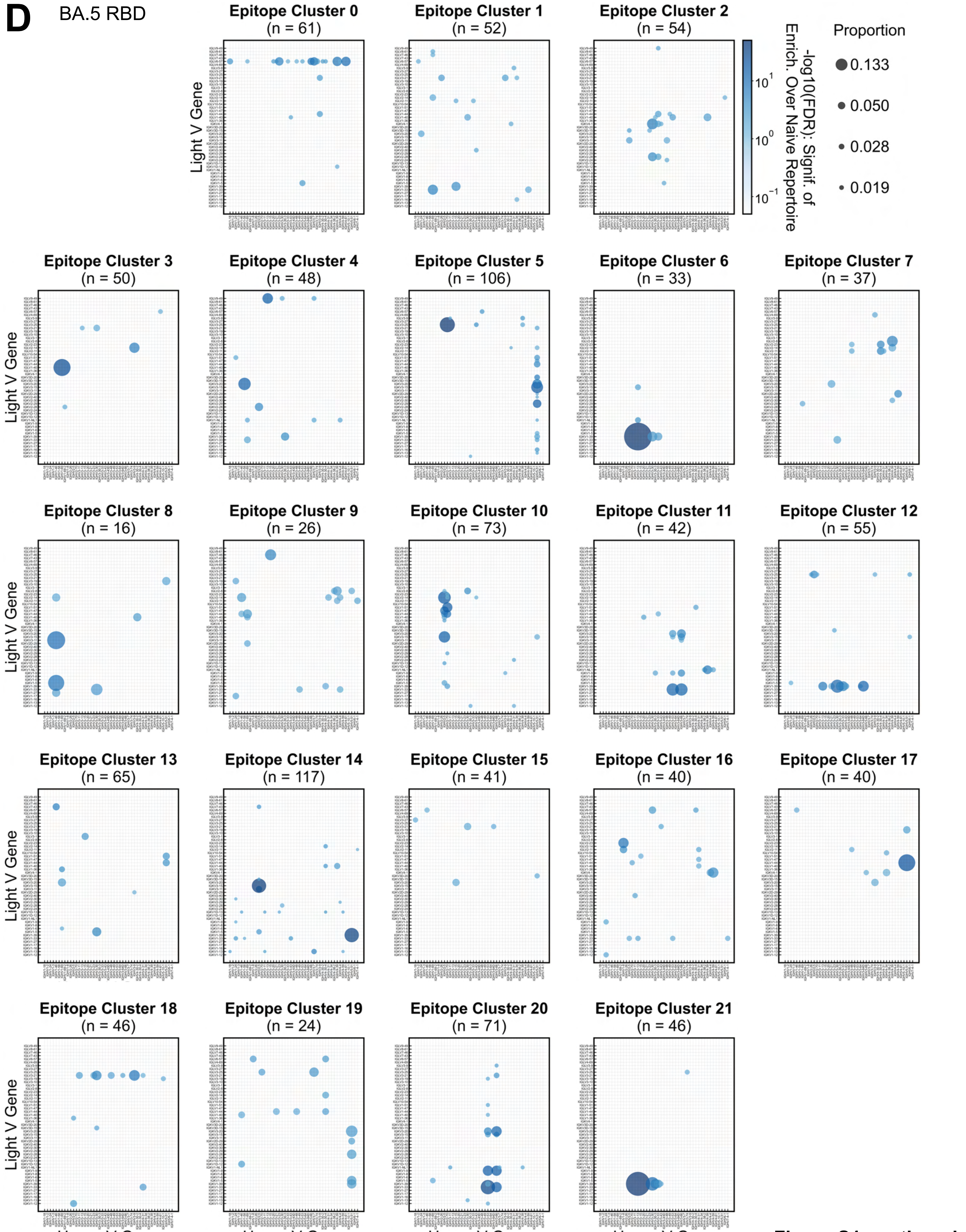


Figure S1 continued

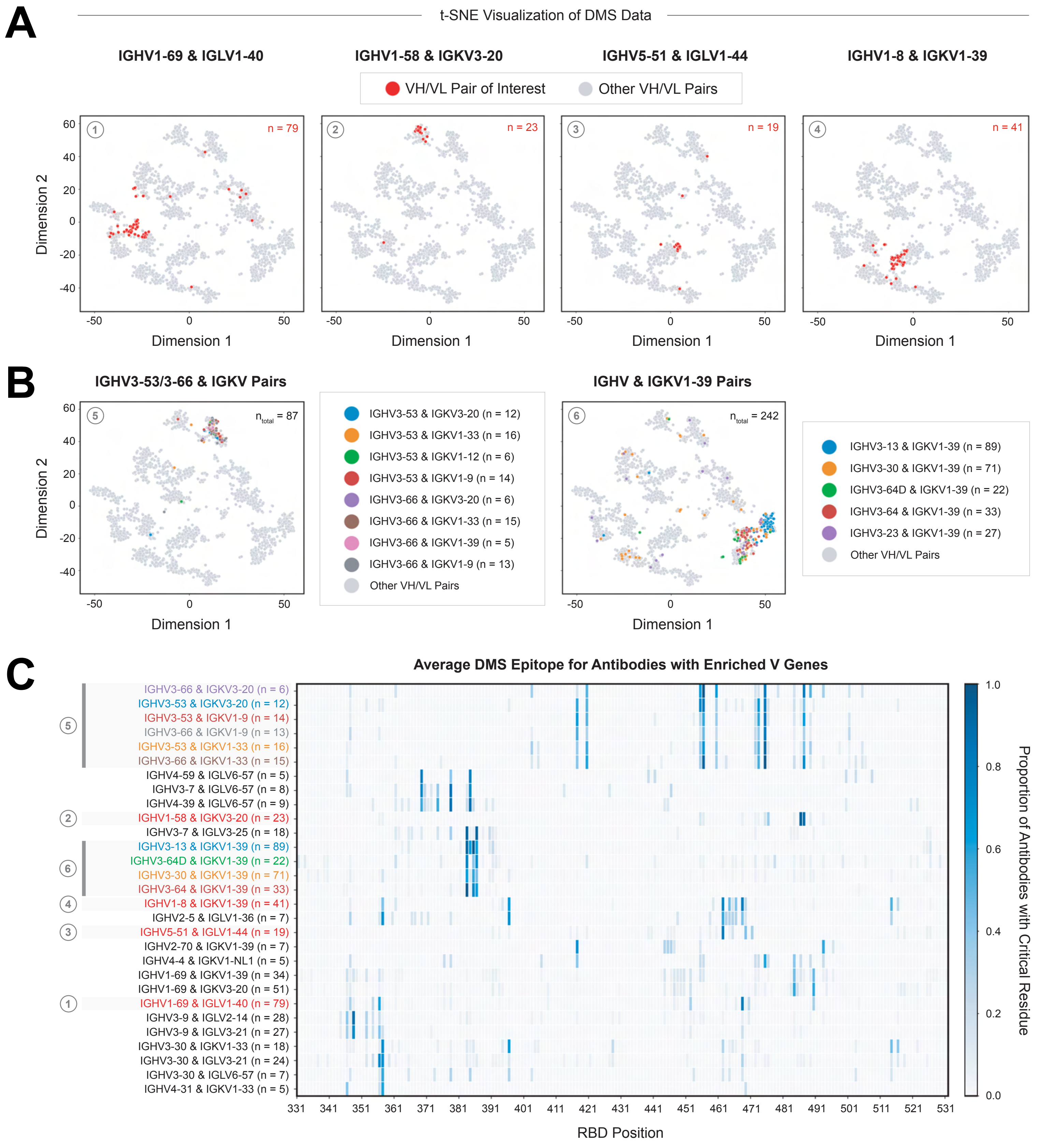
A
t-SNE Visualization of DMS Data
IGHV1-69 & IGLV1-40
IGHV1-58 & IGKV3-20
IGHV5-51 & IGLV1-44
IGHV1-8 & IGKV1-39
VH/VL Pair of Interest
Other VH/VL Pairs
n = 79
n = 23
n = 19
n = 41
Dimension 2
Dimension 1
B
IGHV3-53/3-66 & IGKV Pairs
IGHV & IGKV1-39 Pairs
n total = 87
n total = 242
IGHV3-53 & IGKV3-20 (n = 12)
IGHV3-53 & IGKV1-33 (n = 16)
IGHV3-53 & IGKV1-12 (n = 6)
IGHV3-53 & IGKV1-9 (n = 14)
IGHV3-66 & IGKV3-20 (n = 6)
IGHV3-66 & IGKV1-33 (n = 15)
IGHV3-66 & IGKV1-39 (n = 5)
IGHV3-66 & IGKV1-9 (n = 13)
Other VH/VL Pairs
IGHV3-13 & IGKV1-39 (n = 89)
IGHV3-30 & IGKV1-39 (n = 71)
IGHV3-64D & IGKV1-39 (n = 22)
IGHV3-64 & IGKV1-39 (n = 33)
IGHV3-23 & IGKV1-39 (n = 27)
Other VH/VL Pairs
C
Average DMS Epitope for Antibodies with Enriched V Genes
IGHV3-66 & IGKV3-20 (n = 6)
IGHV3-53 & IGKV3-20 (n = 12)
IGHV3-53 & IGKV1-9 (n = 14)
IGHV3-66 & IGKV1-9 (n = 13)
IGHV3-53 & IGKV1-33 (n = 16)
IGHV3-66 & IGKV1-33 (n = 15)
IGHV4-59 & IGLV6-57 (n = 5)
IGHV3-7 & IGLV6-57 (n = 8)
IGHV4-39 & IGLV6-57 (n = 9)
IGHV1-58 & IGKV3-20 (n = 23)
IGHV3-7 & IGLV3-25 (n = 18)
IGHV3-13 & IGKV1-39 (n = 89)
IGHV3-64D & IGKV1-39 (n = 22)
IGHV3-30 & IGKV1-39 (n = 71)
IGHV3-64 & IGKV1-39 (n = 33)
IGHV1-8 & IGKV1-39 (n = 41)
IGHV2-5 & IGLV1-36 (n = 7)
IGHV5-51 & IGLV1-44 (n = 19)
IGHV2-70 & IGKV1-39 (n = 7)
IGHV4-4 & IGKV1-NL1 (n = 5)
IGHV1-69 & IGKV1-39 (n = 34)
IGHV1-69 & IGKV3-20 (n = 51)
IGHV1-69 & IGLV1-40 (n = 79)
IGHV3-9 & IGLV2-14 (n = 28)
IGHV3-9 & IGLV3-21 (n = 27)
IGHV3-30 & IGKV1-33 (n = 18)
IGHV3-30 & IGLV3-21 (n = 24)
IGHV3-30 & IGLV6-57 (n = 7)
IGHV4-31 & IGKV1-33 (n = 5)
RBD Position
Proportion of Antibodies with Critical Residue

Figure S2

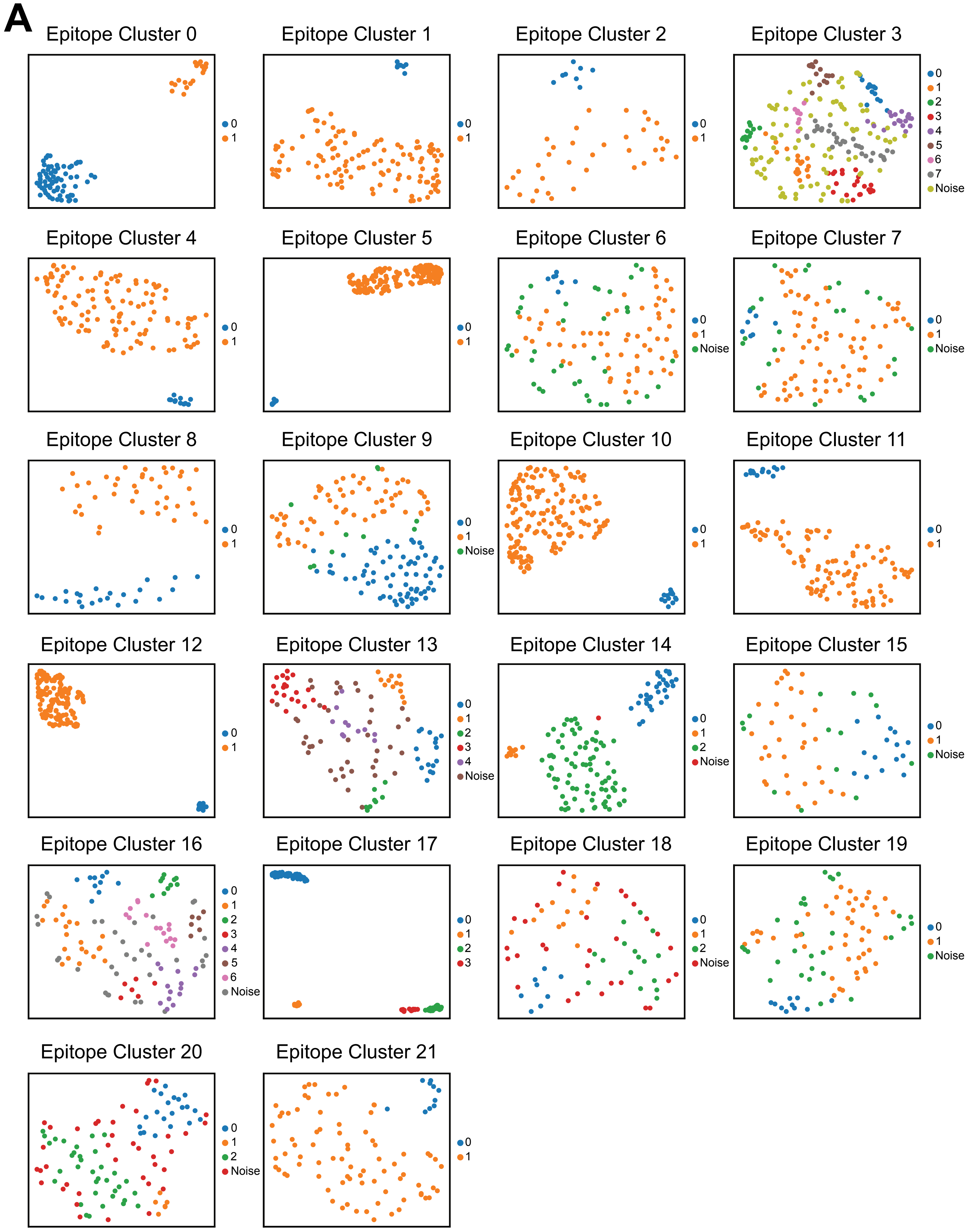

A
Epitope Cluster 0
0
1
Epitope Cluster 1
0
1
Epitope Cluster 2
0
1
Epitope Cluster 3
0
1
2
3
4
5
6
7
Noise
Epitope Cluster 4
0
1
Epitope Cluster 5
0
1
Epitope Cluster 6
0
1
Noise
Epitope Cluster 7
0
1
Noise
Epitope Cluster 8
0
1
Epitope Cluster 9
0
1
Noise
Epitope Cluster 10
0
1
Epitope Cluster 11
0
1
Epitope Cluster 12
0
1
Epitope Cluster 13
0
1
2
3
4
Noise
Epitope Cluster 14
0
1
2
Noise
Epitope Cluster 15
0
1
Noise
Epitope Cluster 16
0
1
2
3
4
5
6
Noise
Epitope Cluster 17
0
1
2
3
Epitope Cluster 18
0
1
2
Noise
Epitope Cluster 19
0
1
Noise
Epitope Cluster 20
0
1
2
Noise
Epitope Cluster 21
0
1


**Figure S3**

**B**

**DMS Epitope Cluster 0** Subcluster 0 (n = 68)

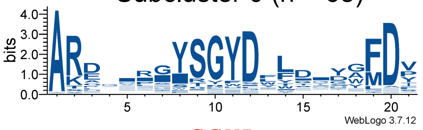

**DMS Epitope Cluster 1** Subcluster 0 (n = 8)

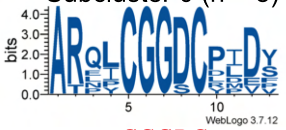

**DMS Epitope Cluster 2** Subcluster 0 (n = 8)

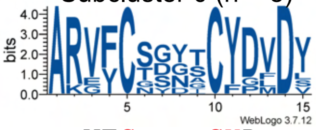

| | Cluster 0 / Subcluster 0 | Cluster 1 / Subcluster 0 | Cluster 2 / Subcluster 0 |
|---|---|---|---|
| Consensus | ------SGYD------ | --CGGDC- | VFC----CYD |
| Gene Segment(s) | IGHD5-12*01 GYSGYDY | IGHD2-21*01 AYCGGDCYS | IGHD2-15*01 GYCSGGSCYS |

**DMS Epitope Cluster 3** Subcluster 0 (n = 13)

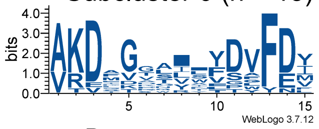

**DMS Epitope Cluster 3** Subcluster 2 (n = 13)

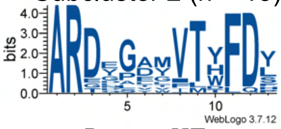

**DMS Epitope Cluster 4** Subcluster 0 (n = 11)

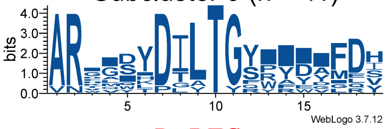

| | Cluster 3 / Subcluster 0 | Cluster 3 / Subcluster 2 | Cluster 4 / Subcluster 0 |
|---|---|---|---|
| Consensus | D--------- | D----VT- | ----D-LTG----- |
| Gene Segment(s) | Undetermined | Undetermined | IGHD3-9*01 YYDILTGYYN |

**DMS Epitope Cluster 5** Subcluster 0 (n = 11)

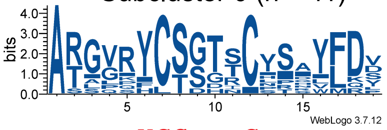

**DMS Epitope Cluster 6** Subcluster 0 (n = 7)

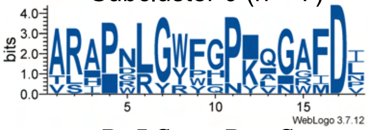

**DMS Epitope Cluster 7** Subcluster 0 (n = 7)

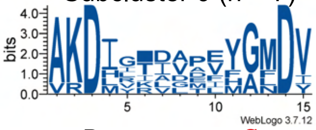

| | Cluster 5 / Subcluster 0 | Cluster 6 / Subcluster 0 | Cluster 7 / Subcluster 0 |
|---|---|---|---|
| Consensus | ---YCS---C---- | -P-LG---P--G- | D--------G |
| Gene Segment(s) | IGHD2-2*01 GYCSSTSCYA<br>IGHD2-15*01 GYCSGGSCYS | Undetermined | IGHJ6*01 YYYYYGMDV... |

**DMS Epitope Cluster 8** Subcluster 0 (n = 23)

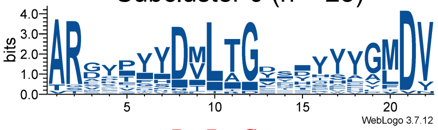

**DMS Epitope Cluster 8** Subcluster 1 (n = 40)

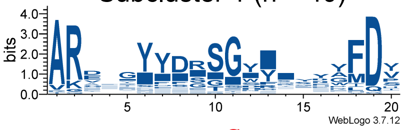

**DMS Epitope Cluster 10** Subcluster 0 (n = 13)

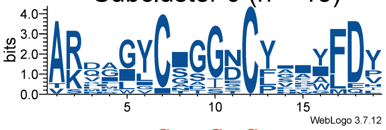

| | Cluster 8 / Subcluster 0 | Cluster 8 / Subcluster 1 | Cluster 10 / Subcluster 0 |
|---|---|---|---|
| Consensus | -----D-L-G------- | ---------G------ | ----C--G-C---- |
| Gene Segment(s) | IGHD3-9*01 YYDILTGYYN | IGHD3-22*01 YYYDSSGYYY | IGHD2-21*01 AYC-GGDCYS<br>IGHD2-15*01 GYCSGGSCYS |

**DMS Epitope Cluster 11** Subcluster 0 (n = 13)

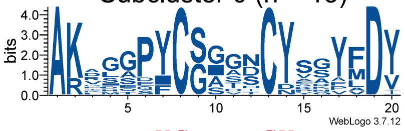

**DMS Epitope Cluster 12** Subcluster 0 (n = 20)

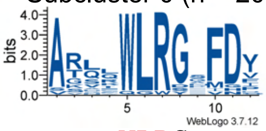

**DMS Epitope Cluster 13** Subcluster 1 (n = 12)

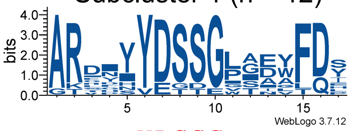

| | Cluster 11 / Subcluster 0 | Cluster 12 / Subcluster 0 | Cluster 13 / Subcluster 1 |
|---|---|---|---|
| Consensus | ----YC----CY--- | --WLRG- | ---YDSSG---- |
| Gene Segment(s) | IGHD2-2*01 GYCSSTSCYA<br>IGHD2-15*01 GYCSGGSCYS<br>IGHD2-21*01 AYCG-GDCYS | IGHD5-12*01 WI*WLRL<br>IGHD5-18*02 **WLRL | IGHD3-22*01 YYYDSSGYYY |

**DMS Epitope Cluster 13** Subcluster 2 (n = 8)

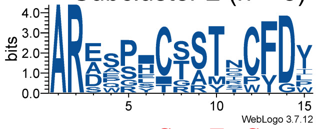

**DMS Epitope Cluster 13** Subcluster 3 (n = 20)

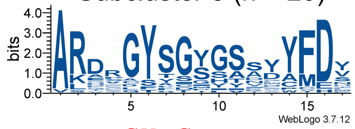

**DMS Epitope Cluster 14** Subcluster 0 (n = 31)

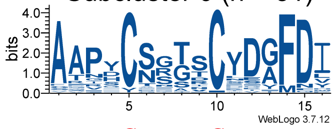

| | Cluster 13 / Subcluster 2 | Cluster 13 / Subcluster 3 | Cluster 14 / Subcluster 0 |
|---|---|---|---|
| Consensus | ----C--T-C | --GY-G------ | --C----C--- |
| Gene Segment(s) | IGHD2-2*01 GYCSSTSCYA | IGHD5-12*01 GYSGYDY | IGHD2-2*01 GYCSSTSCYA<br>IGHD2-15*01 GYCSGGSCYS |

**DMS Epitope Cluster 14** Subcluster 1 (n = 9)

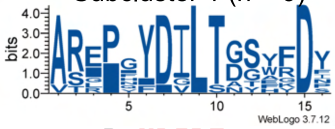

**DMS Epitope Cluster 16** Subcluster 0 (n = 10)

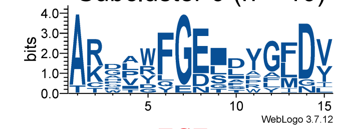

**DMS Epitope Cluster 16** Subcluster 2 (n = 10)

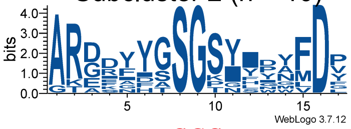

| | Cluster 14 / Subcluster 1 | Cluster 16 / Subcluster 0 | Cluster 16 / Subcluster 2 |
|---|---|---|---|
| Consensus | -P-YDILT--- | ---FGE---- | -----SGS---- |
| Gene Segment(s) | IGHD3-9*01 YYDILTGYYN | IGHD3-10*01 VLLWFGELL* | IGHD3-10*01 YYYGSGSYYN |

**Figure S3 continued**

**DMS Epitope Cluster 16**
Subcluster 3 (n = 6)

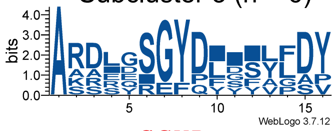

Consensus ---SGYD----
Gene Segment(s) IGHD5-12*01 GYSGYDY

**DMS Epitope Cluster 16**
Subcluster 5 (n = 6)

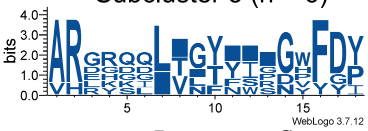

Consensus ----L------G-
Gene Segment(s) Undetermined

**DMS Epitope Cluster 16**
Subcluster 6 (n = 16)

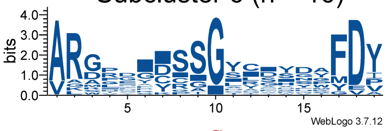

Consensus --------G------
Gene Segment(s)
IGHD3-22*01 YYYDSSGYYY
IGHD6-19*01 GYSSGWY
IGHD6-25*01 GYSSGY
IGHD2-15*01 GYCSGGSCYS

**DMS Epitope Cluster 17**
Subcluster 1 (n = 14)

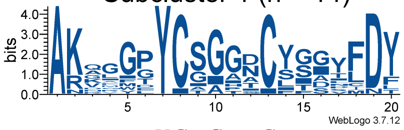

Consensus ----YC-G--C----
Gene Segment(s) IGHD2-15*01 GYCSGGSCYS

**DMS Epitope Cluster 17**
Subcluster 2 (n = 30)

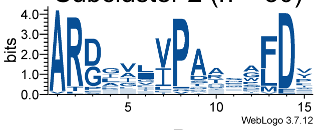

Consensus -----P----
Gene Segment(s) Undetermined

**DMS Epitope Cluster 17**
Subcluster 3 (n = 15)

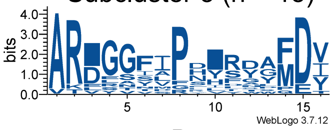

Consensus -----P-----
Gene Segment(s) Undetermined

**DMS Epitope Cluster 18**
Subcluster 0 (n = 9)

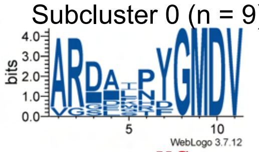

Consensus ----YG
Gene Segment(s) IGHJ6*01 YYYYYGMDV...

**DMS Epitope Cluster 19**
Subcluster 0 (n = 10)

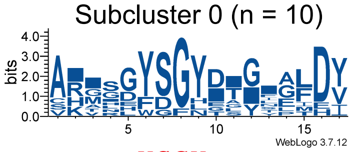

Consensus ---YSGY-----
Gene Segment(s) IGHD5-12*01 GYSGYDY

**DMS Epitope Cluster 20**
Subcluster 1 (n = 6)

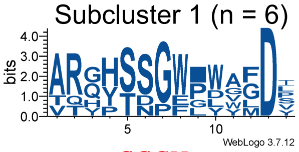

Consensus --SSGW---
Gene Segment(s) IGHD6-19*01 GYSSGWY

**DMS Epitope Cluster 21**
Subcluster 0 (n = 10)

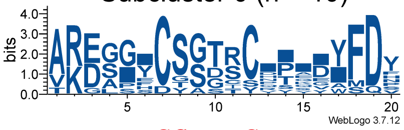

Consensus ----CS---C-----
Gene Segment(s)
IGHD2-15*01 GYCSGGSCYS
IGHD2-2*01 GYCSSTSCYA

**Figure S3 continued**

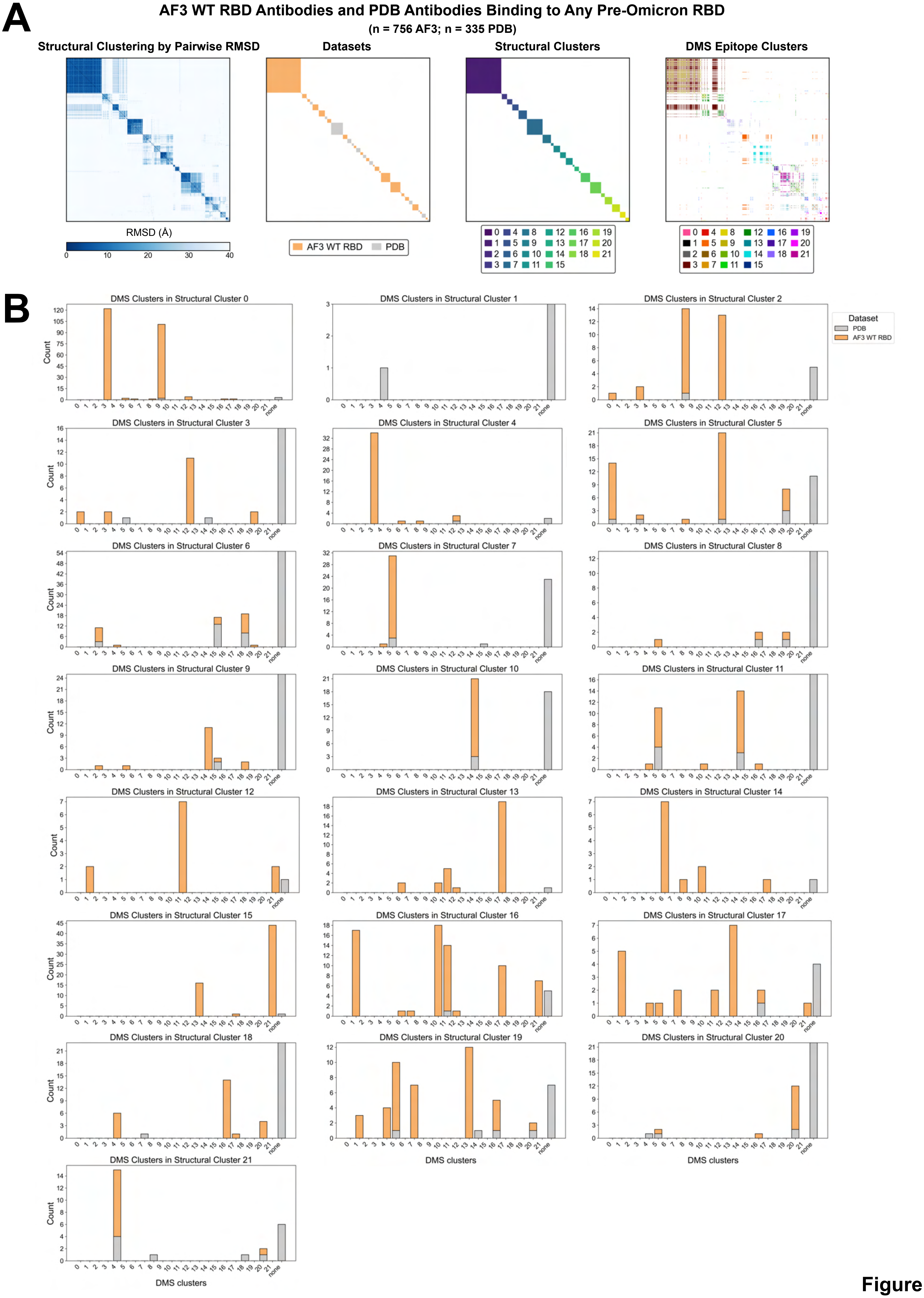
A
AF3 WT RBD Antibodies and PDB Antibodies Binding to Any Pre-Omicron RBD
(n = 756 AF3; n = 335 PDB)
Structural Clustering by Pairwise RMSD
Datasets
Structural Clusters
DMS Epitope Clusters
RMSD (Å)
0 10 20 30 40
AF3 WT RBD
PDB
0 1 2 3 4 5 6 7 8 9 10 11 12 13 14 15 16 17 18 19 20 21
B
DMS Clusters in Structural Cluster 0
DMS Clusters in Structural Cluster 1
DMS Clusters in Structural Cluster 2
DMS Clusters in Structural Cluster 3
DMS Clusters in Structural Cluster 4
DMS Clusters in Structural Cluster 5
DMS Clusters in Structural Cluster 6
DMS Clusters in Structural Cluster 7
DMS Clusters in Structural Cluster 8
DMS Clusters in Structural Cluster 9
DMS Clusters in Structural Cluster 10
DMS Clusters in Structural Cluster 11
DMS Clusters in Structural Cluster 12
DMS Clusters in Structural Cluster 13
DMS Clusters in Structural Cluster 14
DMS Clusters in Structural Cluster 15
DMS Clusters in Structural Cluster 16
DMS Clusters in Structural Cluster 17
DMS Clusters in Structural Cluster 18
DMS Clusters in Structural Cluster 19
DMS Clusters in Structural Cluster 20
DMS Clusters in Structural Cluster 21
Dataset
PDB
AF3 WT RBD
Count
DMS clusters
none

Figure S4

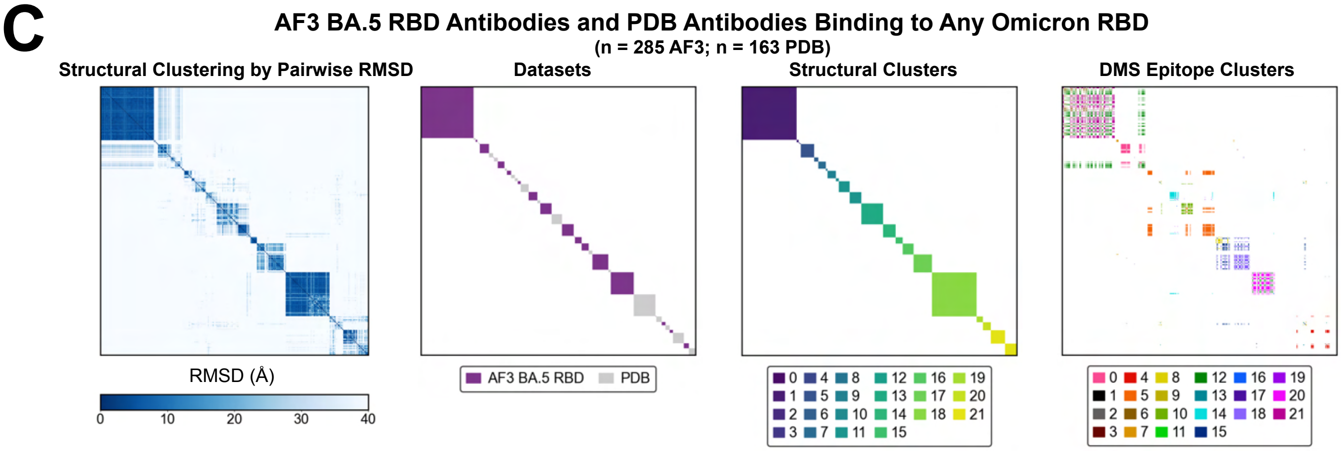


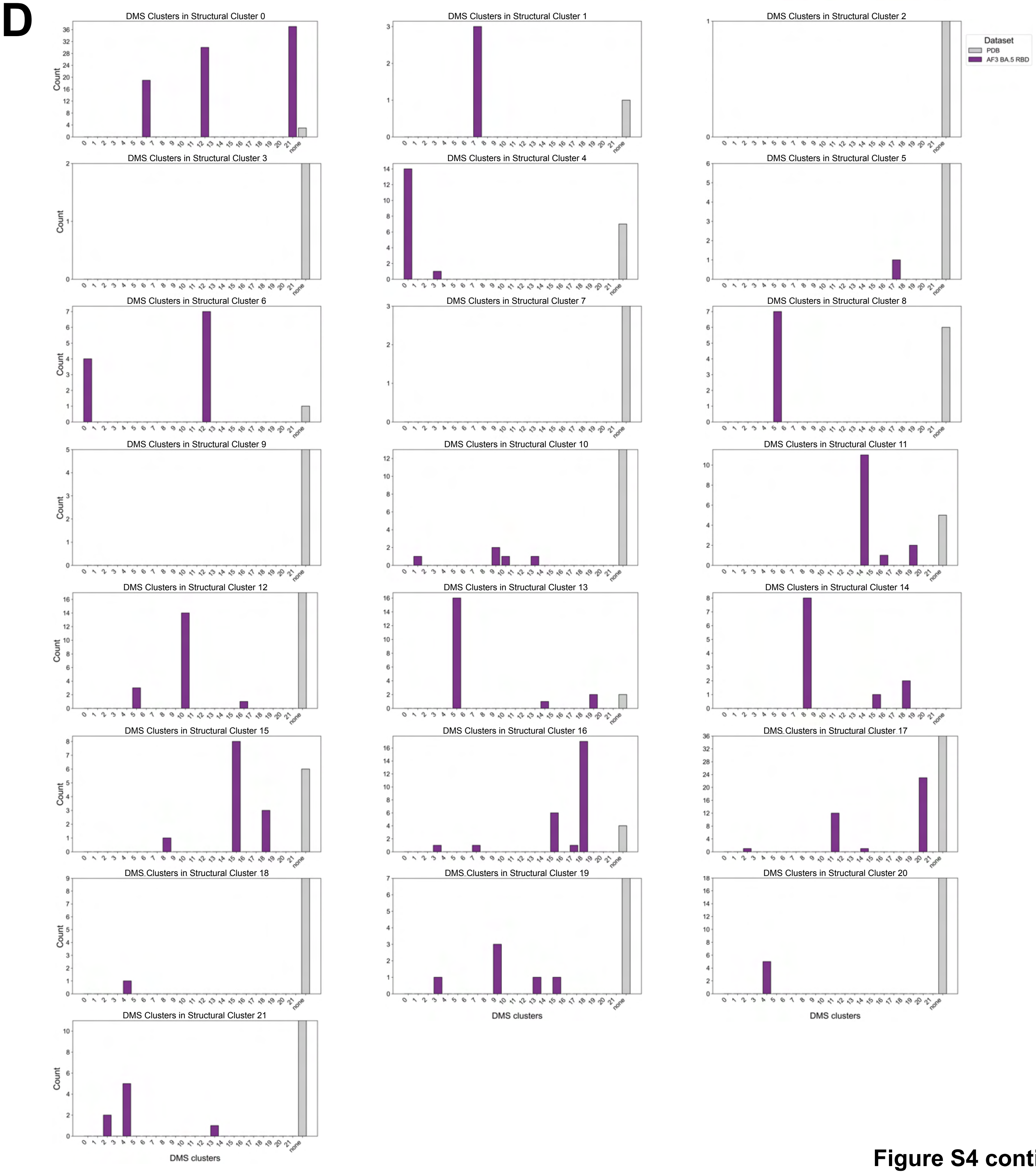


**Figure S4 continued**

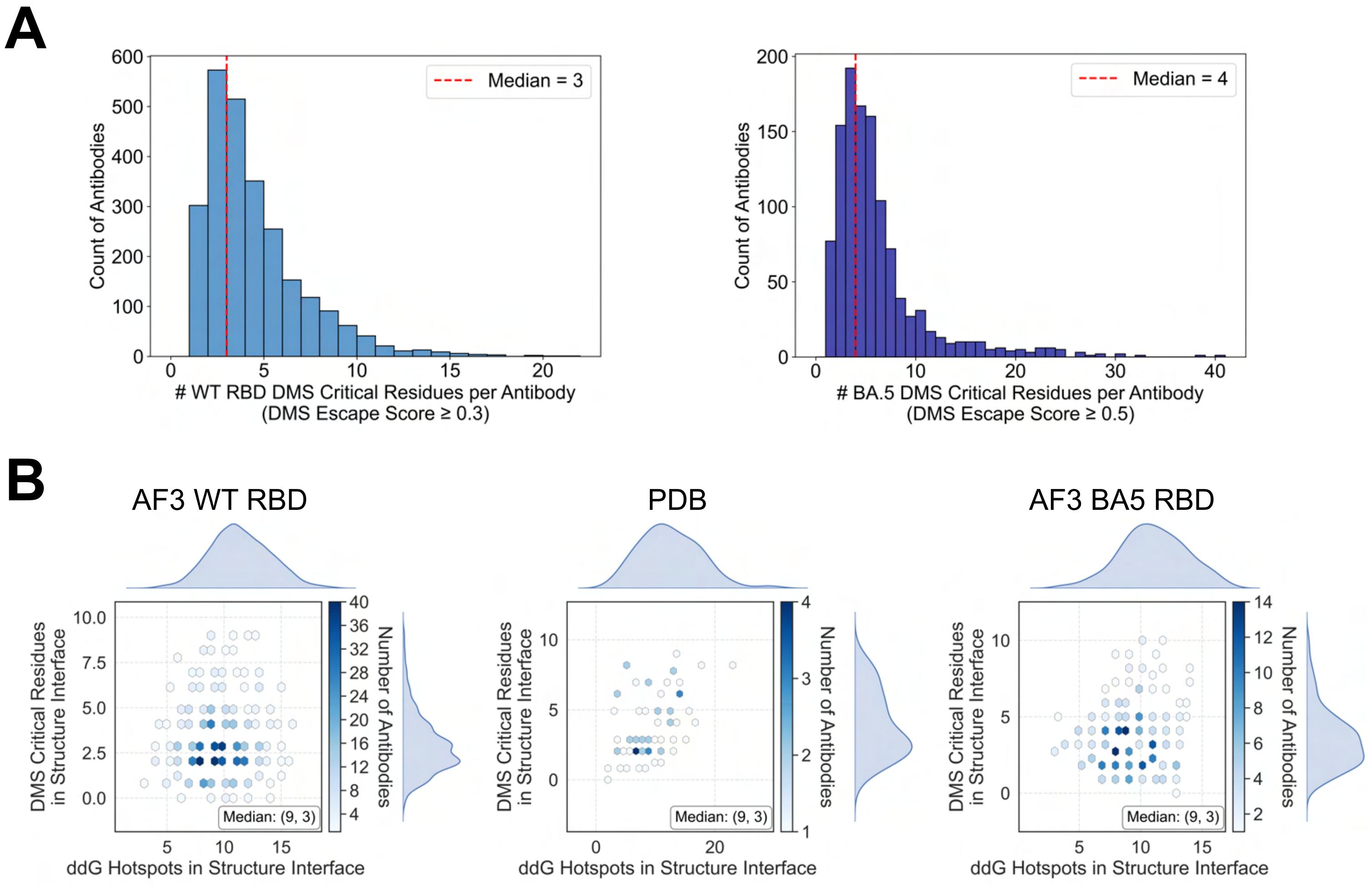
A
Count of Antibodies
Median = 3
### WT RBD DMS Critical Residues per Antibody
(DMS Escape Score ≥ 0.3)
Median = 4
### BA.5 DMS Critical Residues per Antibody
(DMS Escape Score ≥ 0.5)
B
AF3 WT RBD
PDB
AF3 BA5 RBD
DMS Critical Residues in Structure Interface
ddG Hotspots in Structure Interface
Number of Antibodies
Median: (9, 3)

Figure S5

C

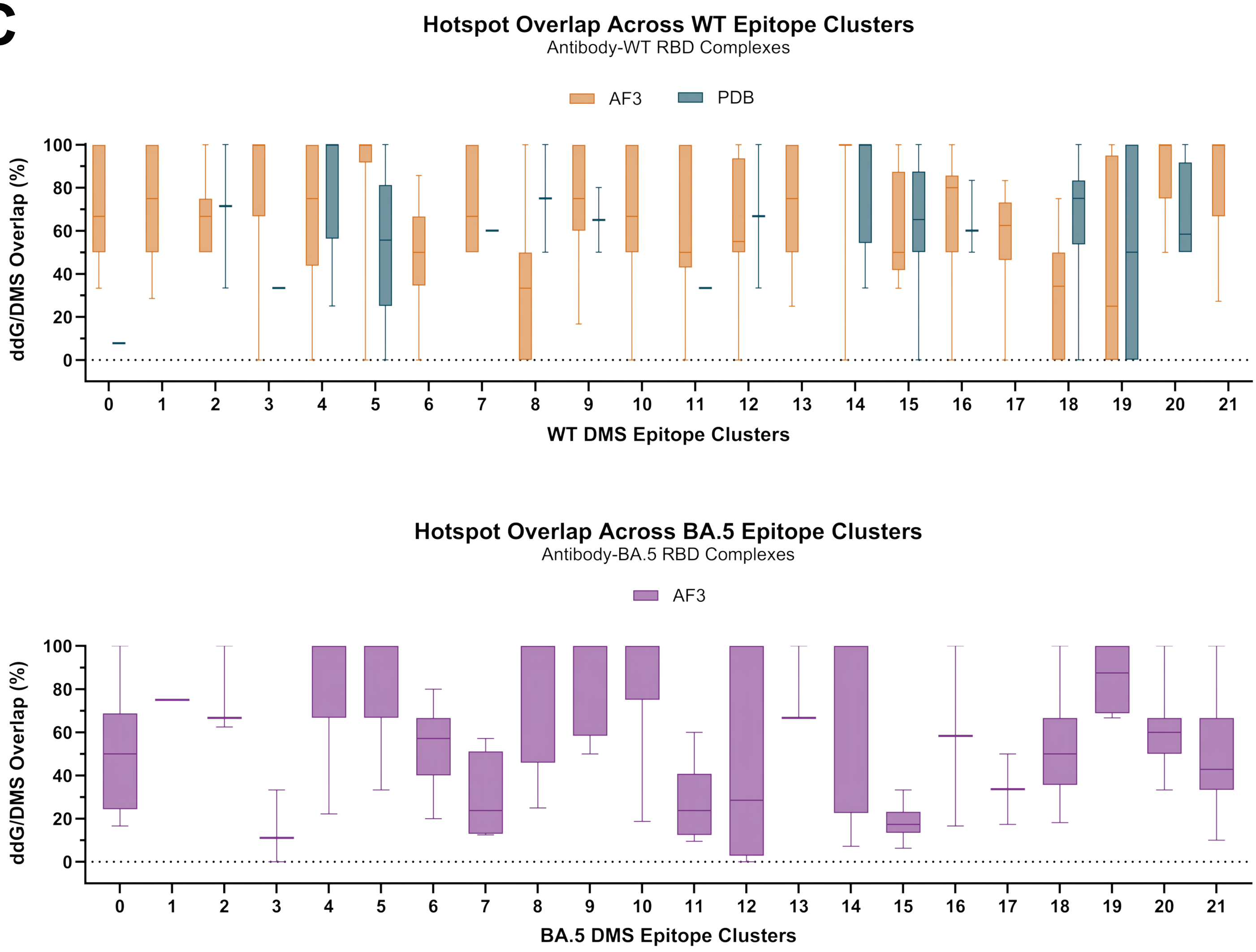


Figure S5 continued

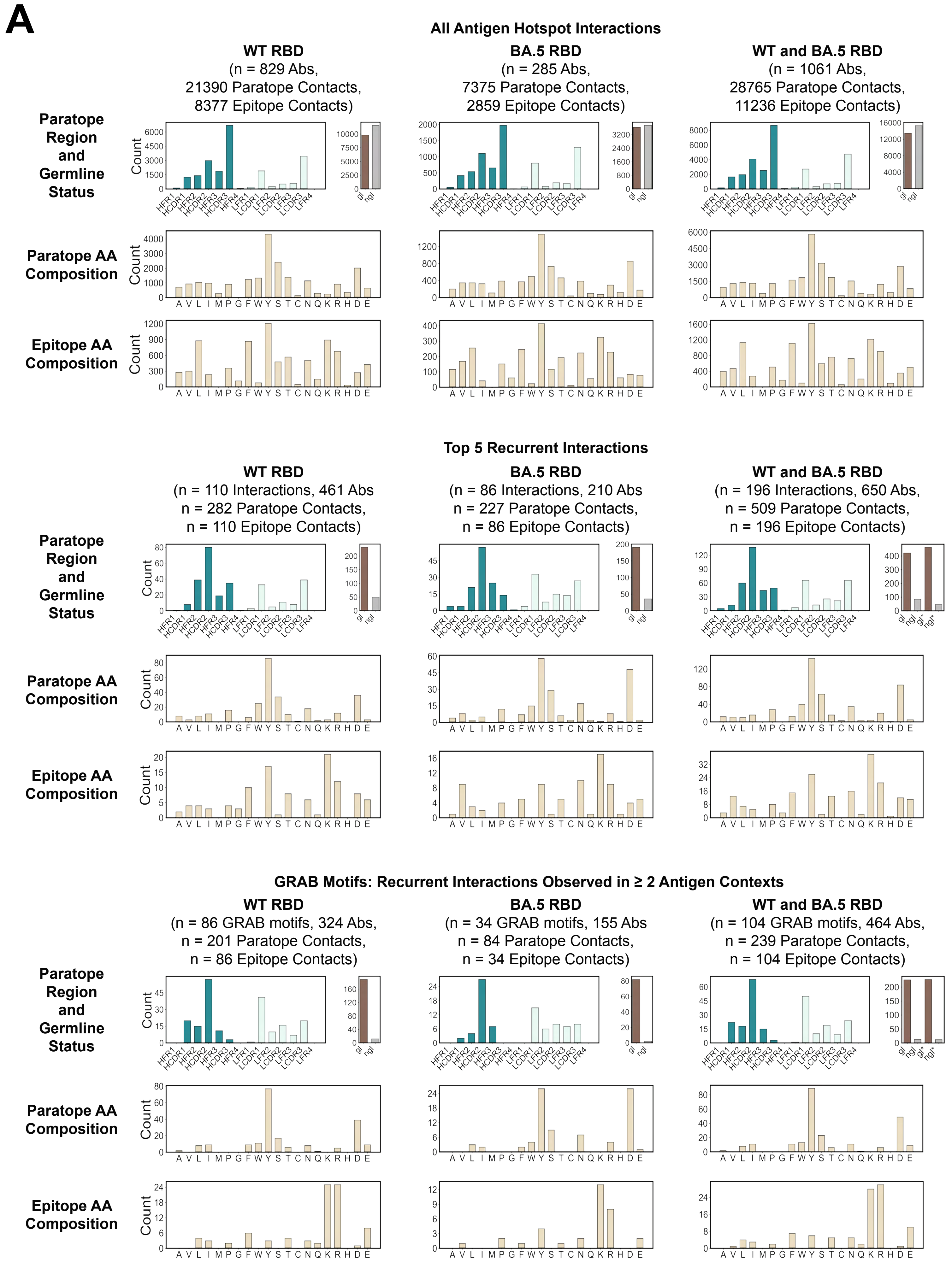
A
All Antigen Hotspot Interactions
WT RBD
(n = 829 Abs,
21390 Paratope Contacts,
8377 Epitope Contacts)
BA.5 RBD
(n = 285 Abs,
7375 Paratope Contacts,
2859 Epitope Contacts)
WT and BA.5 RBD
(n = 1061 Abs,
28765 Paratope Contacts,
11236 Epitope Contacts)
Paratope Region and Germline Status
Paratope AA Composition
Epitope AA Composition
Count
Top 5 Recurrent Interactions
WT RBD
(n = 110 Interactions, 461 Abs
n = 282 Paratope Contacts,
n = 110 Epitope Contacts)
BA.5 RBD
(n = 86 Interactions, 210 Abs
n = 227 Paratope Contacts,
n = 86 Epitope Contacts)
WT and BA.5 RBD
(n = 196 Interactions, 650 Abs,
n = 509 Paratope Contacts,
n = 196 Epitope Contacts)
GRAB Motifs: Recurrent Interactions Observed in ≥ 2 Antigen Contexts
WT RBD
(n = 86 GRAB motifs, 324 Abs,
n = 201 Paratope Contacts,
n = 86 Epitope Contacts)
BA.5 RBD
(n = 34 GRAB motifs, 155 Abs
n = 84 Paratope Contacts,
n = 34 Epitope Contacts)
WT and BA.5 RBD
(n = 104 GRAB motifs, 464 Abs,
n = 239 Paratope Contacts,
n = 104 Epitope Contacts)

Figure S6

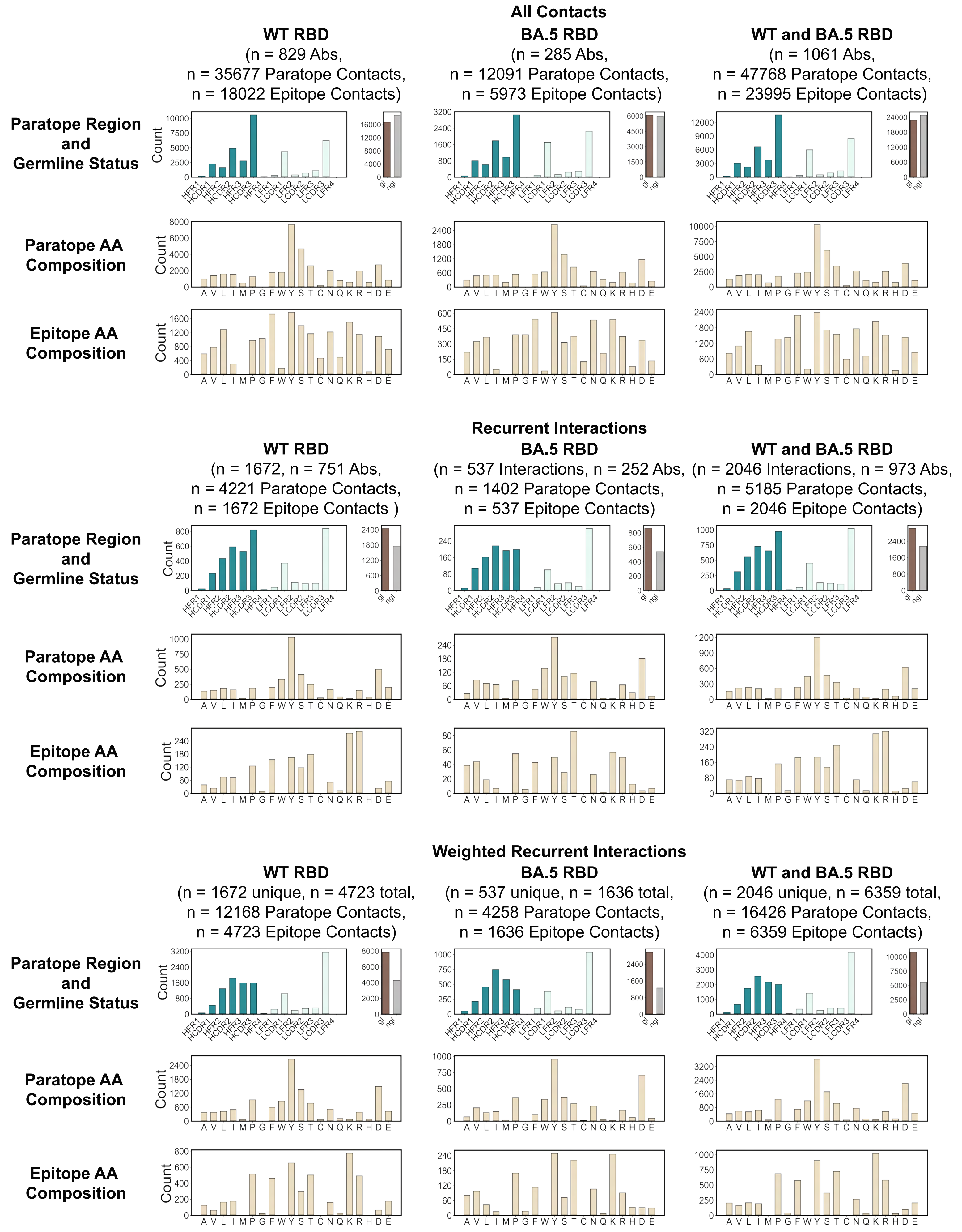


**Figure S6 continued**

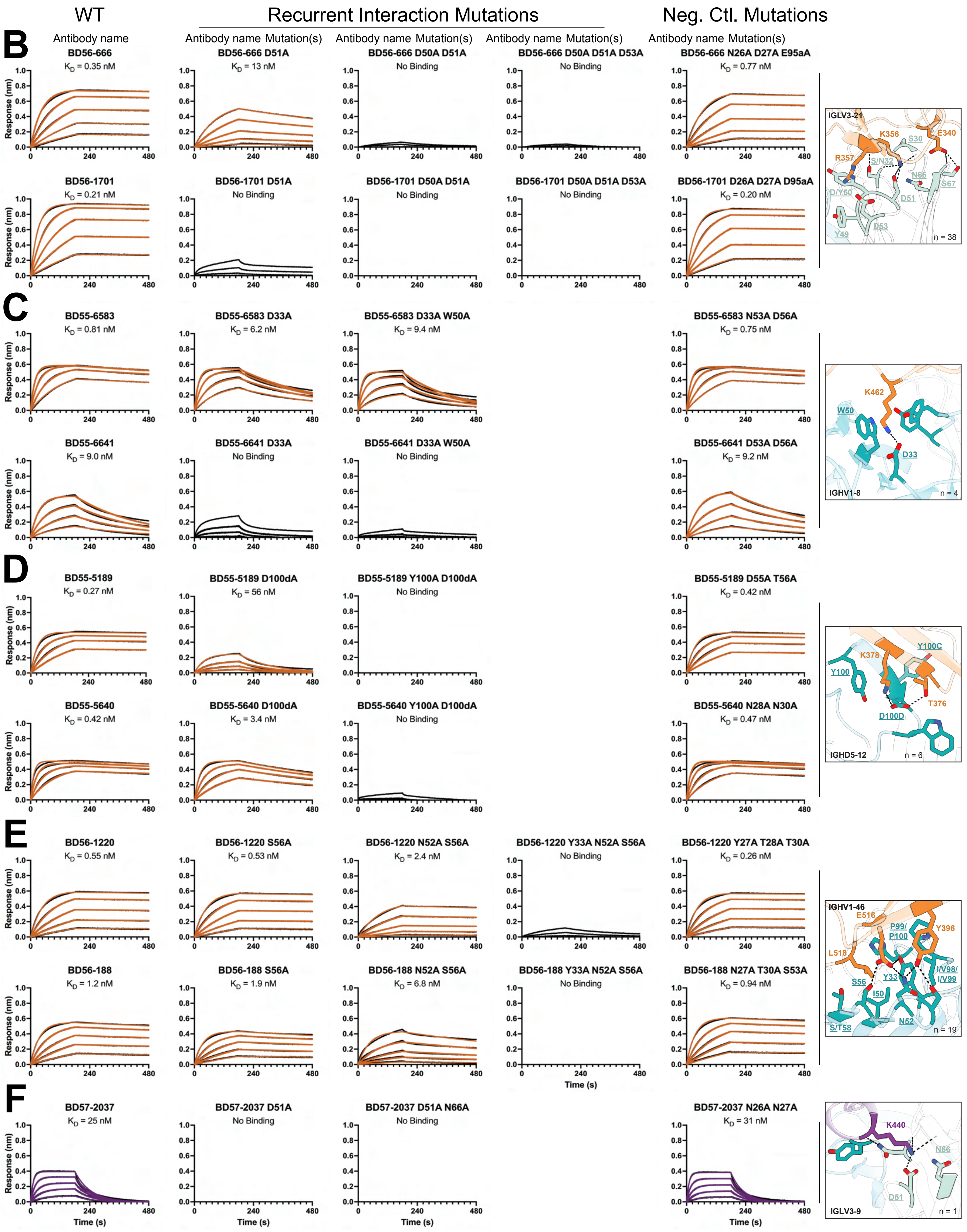


**Figure S6 continued**

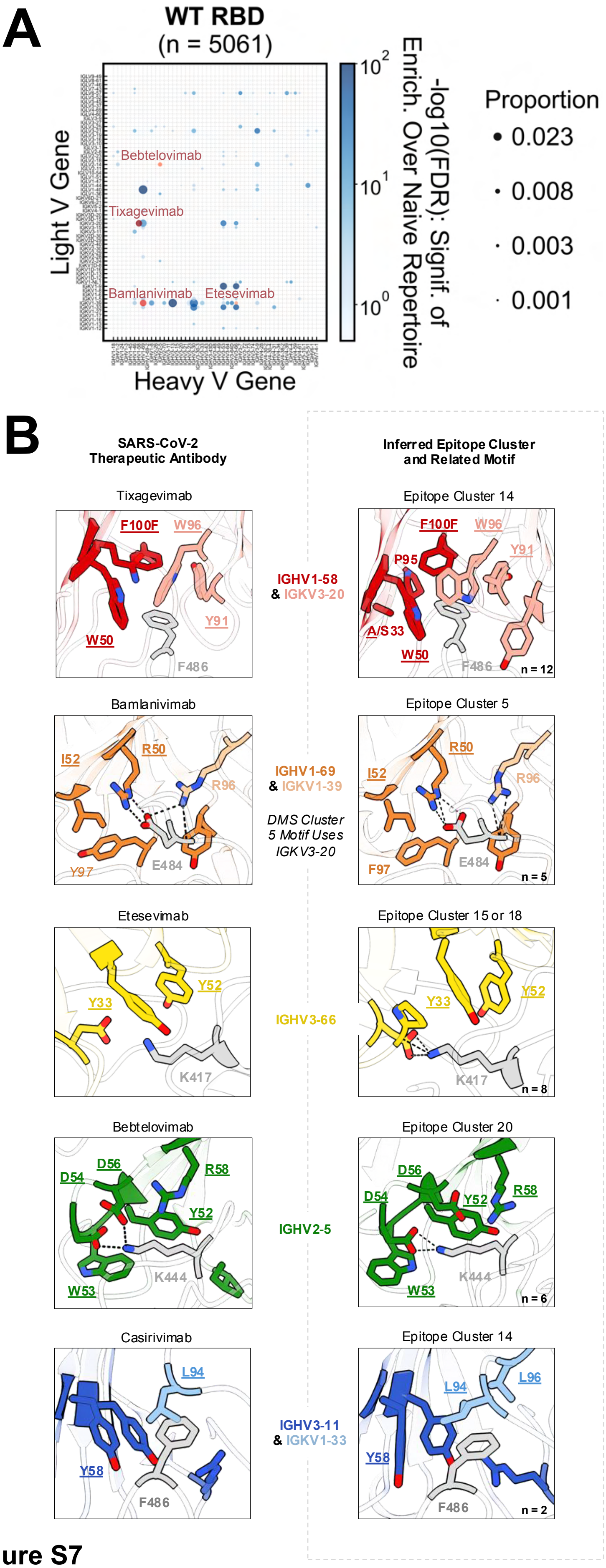
A
WT RBD
(n = 5061)
Light V Gene
Heavy V Gene
Bebtelovimab
Tixagevimab
Bamlanivimab
Etesevimab
-log10(FDR): Signif. of Enrich. Over Naive Repertoire
Proportion
0.023
0.008
0.003
0.001
B
SARS-CoV-2 Therapeutic Antibody
Inferred Epitope Cluster and Related Motif
Tixagevimab
Epitope Cluster 14
IGHV1-58 & IGKV3-20
F100F
W96
Y91
W50
F486
P95
A/S33
n = 12
Bamlanivimab
Epitope Cluster 5
IGHV1-69 & IGKV1-39
DMS Cluster 5 Motif Uses IGKV3-20
I52
R50
R96
Y97
E484
F97
n = 5
Etesevimab
Epitope Cluster 15 or 18
IGHV3-66
Y33
Y52
K417
n = 8
Bebtelovimab
Epitope Cluster 20
IGHV2-5
D54
D56
R58
Y52
W53
K444
n = 6
Casirivimab
Epitope Cluster 14
IGHV3-11 & IGKV1-33
L94
L96
Y58
F486
n = 2

**Figure S7**

**C**

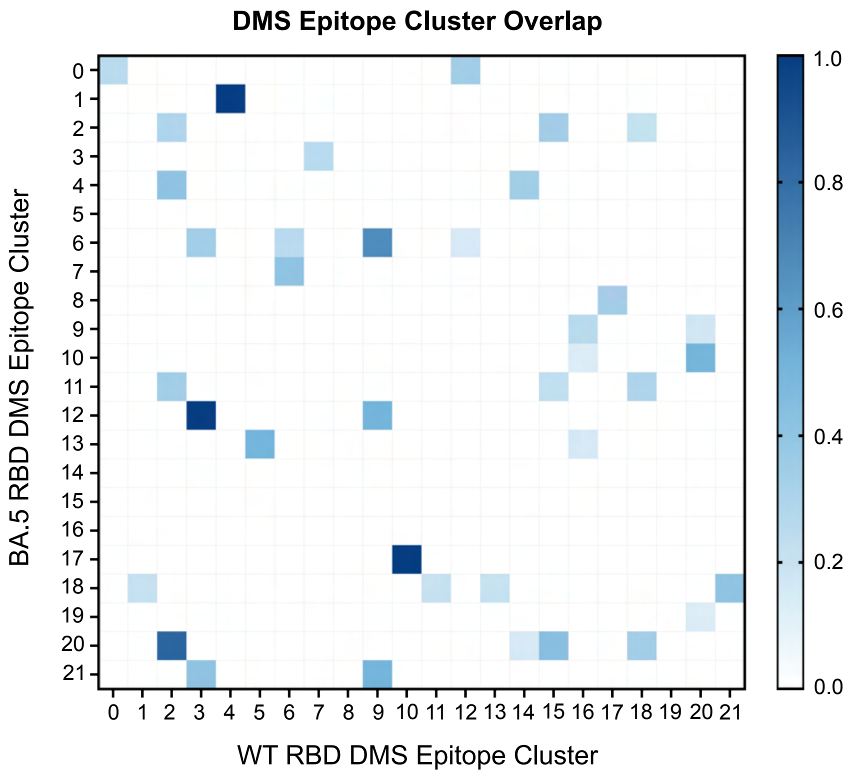


##### WT RBD DMS Epitope Clusters

| Clust. | Critical Residue(s) |
|---|---|
| 0 | 376, 378, 408 |
| 1 | 357, 468 |
| 2 | 455, 456, 475, 487, 489 |
| 3 | 383, 386 |
| 4 | 346 |
| 5 | 484, 490 |
| 6 | 391, 393, 518, 525 |
| 7 | 348, 352 |
| 8 | 381, 413 |
| 9 | 383, 384, 385, 386 |
| 10 | 462 |
| 11 | 357, 396 |
| 12 | 378, 384 |
| 13 | 356, 468 |
| 14 | 486, 487 |
| 15 | 417, 455, 456, 475 |
| 16 | 444, 447, 448, 449, 450, 452, 490 |
| 17 | 396, 514, 516 |
| 18 | 456, 475 |
| 19 | 503, 504 |
| 20 | 444, 445, 446, 447 |
| 21 | 356, 357, 468 |

##### BA.5 RBD DMS Epitope Clusters

| Clust. | Critical Residue(s) |
|---|---|
| 0 | 377, 378 |
| 1 | 346 |
| 2 | 403, 455, 456, 505 |
| 3 | 336, 348, 361 |
| 4 | 487, 489 |
| 5 | 439, 440 |
| 6 | 383, 384, 385, 386, 391, 525 |
| 7 | 391, 524, 525 |
| 8 | 516 |
| 9 | 447, 449, 498 |
| 10 | 444, 445 |
| 11 | 420, 421, 456, 473, 475, 480, 489 |
| 12 | 383, 386 |
| 13 | 490 |
| 14 | 505 |
| 15 | 336, 353, 361, 399, 454, 466, 495, 497 |
| 16 | 498 |
| 17 | 462 |
| 18 | 336, 356, 357, 361 |
| 19 | 439, 445, 498, 499, 500 |
| 20 | 455, 456, 460, 475, 487, 489 |
| 21 | 336, 361, 383, 385, 386 |

**Figure S7 continued**

D

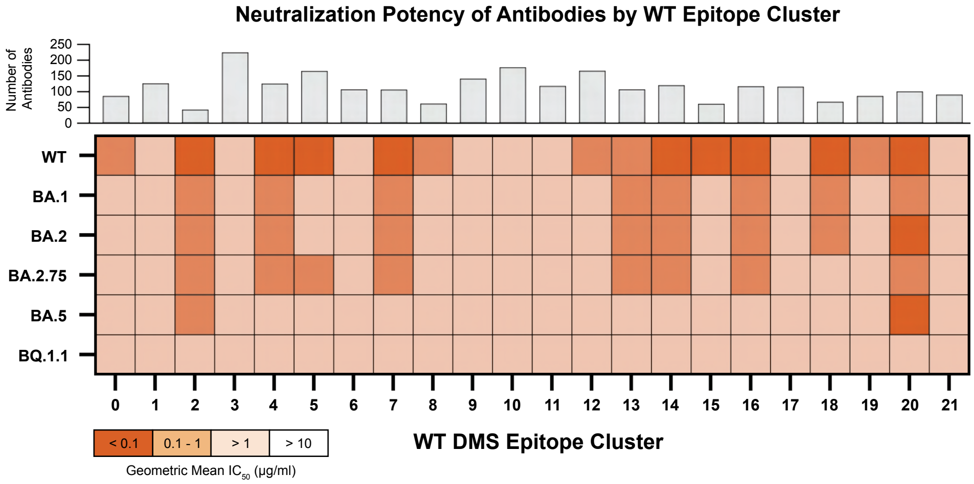


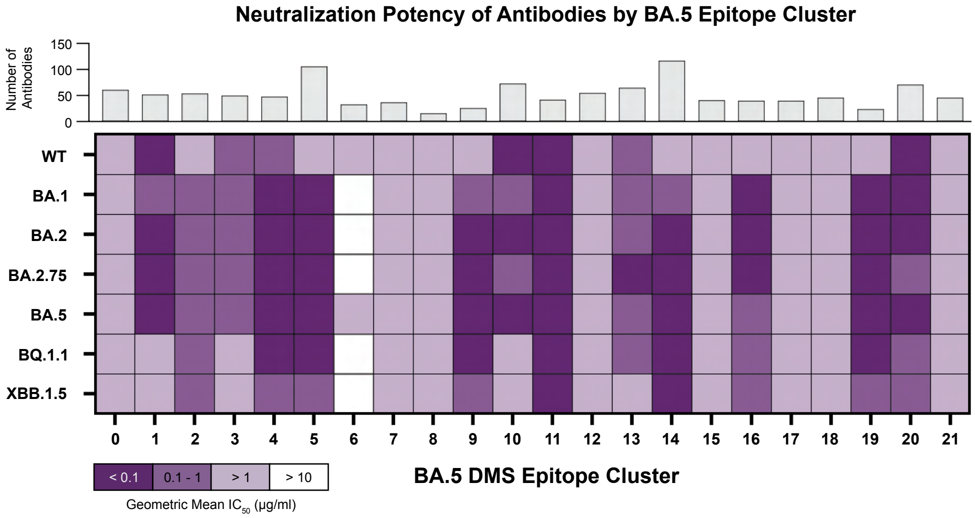


**Figure S7 continued**
